# Simultaneous Capillary Electrophoresis - Mass Spectrometry profiling of (p)ppGpp and dinucleoside polyphosphates reveals oscillatory (p)ppGpp dynamics and a potential synchronisation between alarmones

**DOI:** 10.64898/2026.01.08.698462

**Authors:** Isabel Prucker, Martin Milanov, Patrick Moser, Christoph Popp, Thomas M. Haas, Danye Qiu, Hans-Georg Koch, Henning J. Jessen

## Abstract

To survive stressful conditions, bacteria enter the stringent response mediated by the magic spot nucleotides (MSN) guanosine 3’5’-bispyrophosphate (ppGpp) and guanosine 3’-diphosphate 5’-triphosphate (pppGpp). Another group of so-called alarmones elevated under stress are dinucleoside polyphosphates (Np_n_Ns), such as diadenosine triphosphate (Ap_3_A). How these two groups of alarmones intersect with one another is poorly understood. Here, we present a sensitive method using capillary electrophoresis coupled to mass spectrometry (CE-MS) with heavy isotope labeled internal references to separate and quantify magic spot nucleotides in parallel with Np_n_Ns. This approach uncovered oscillatory fluctuations in MSN and some Np_n_N during exponential growth, suggesting more dynamic regulation of the stringent response than previously thought. In addition, up to eleven Np_n_Ns were detected after prolonged growth. Np_n_N levels were also quantified under various amino acid starvation conditions, revealing a pronounced accumulation in late stationary phase. In summary, our parallel assignment and quantification of highly charged signaling molecules in bacteria under stress conditions paves the way to better study and understand the cross-talk between different alarmones.

## Introduction

Bacteria are well known to adapt to different stressful conditions like antibiotic treatments^[1]^, oxidative stress^[2]^ or nutrient limitation, e.g. of amino acids^[3]^, and carbon or fatty acid deficiency^[4,5]^. This adaptation is called the “stringent response” and is mainly regulated by the magic spot nucleotides (MSN) guanosine 3’5’-bispyrophosphate (ppGpp) and guanosine 3’-diphosphate 5’-triphosphate (pppGpp), hereafter abbreviated as (p)ppGpp (figure 1A)^[6]^. During the bacterial stringent response, the alarmones (p)ppGpp accumulate in response to nutrient deprivation (especially amino acid starvation). The elevated (p)ppGpp direct RNA polymerase to different stress-responsive promoters^[5,7–9]^. This induces a metabolic adaptation program, which upregulates proteins involved in stress survival, slowing down metabolism and growth, but increasing persistence^[9,10]^. The “stringent response” shifts transcription away from ribosomal RNA synthesis, which is rate-limiting for the energy consuming ribosome biogenesis^[11]^. In *E. coli*, (p)ppGpp are synthesized by RelA directly from GDP or GTP and to a lesser extent by SpoT^[12]^. While SpoT has both a synthetase and a phosphatase domain to reduce (p)ppGpp after stress^[13]^, RelA is only able to synthesize (p)ppGpp^[14]^.

**Figure 1.**
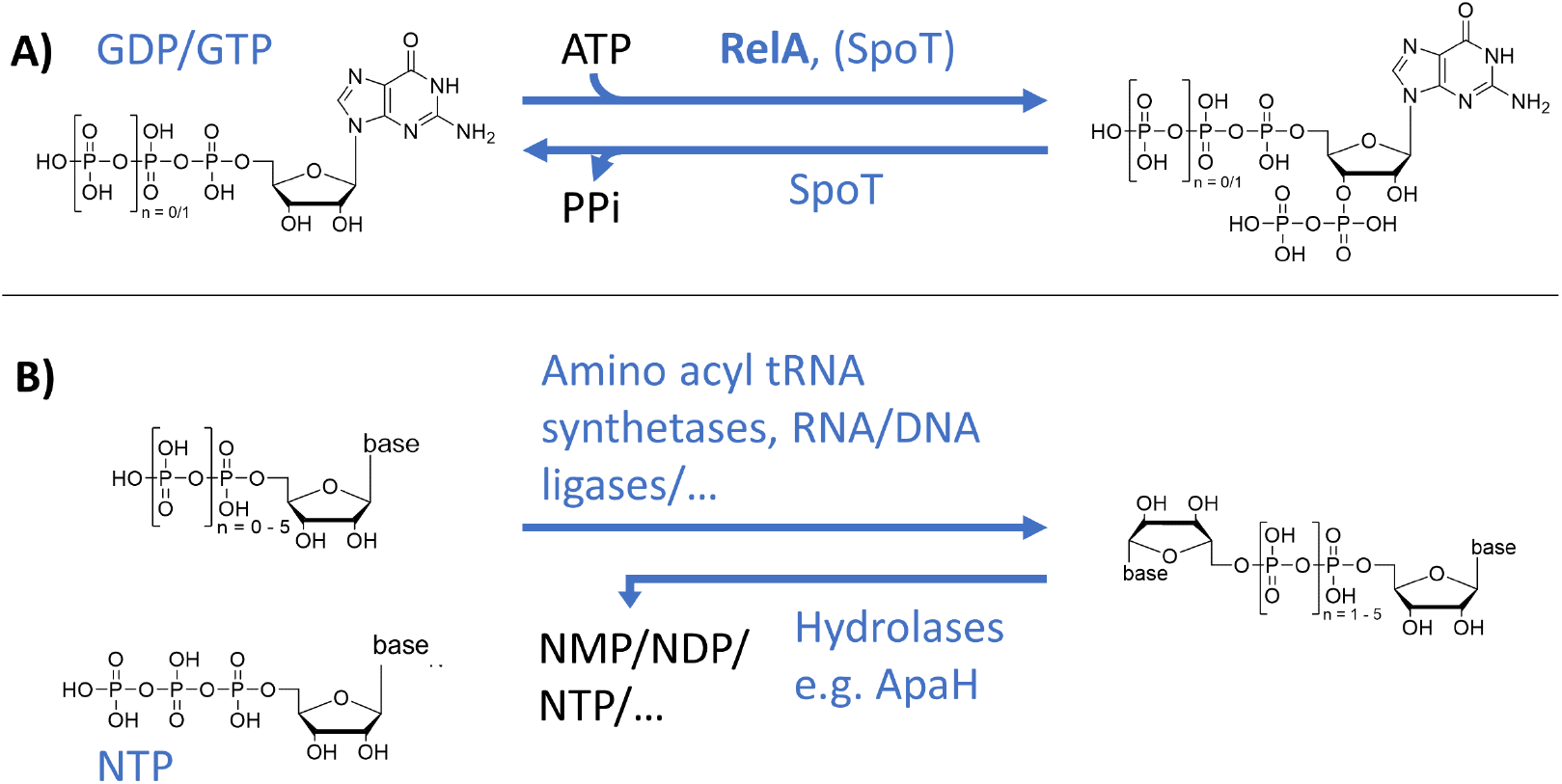
A) Biosynthesis of magic spot nucleotides ppGpp (n = 0) and pppGpp (n = 1). The main synthetase in *E. coli* is RelA. SpoT is also able to synthesize and hydrolyze MSN. B) Synthesis of dinucleoside polyphosphates in bacteria. Various enzymes (e.g. aminoacyl tRNA synthetases^[33]^, firefly luciferase^[43]^ or RNA^[37]^ and DNA^[44]^ ligases) are involved in the biosynthesis. Dinucleoside polyphosphates like Up4U are rare examples of adenine-free variants^[32]^. Hydrolysis is carried out by NUDIX enzymes or ApaH in *E. coli*. Abbreviations: ATP: adenosine 5’-triphosphate, GDP: guanosine 5’-diphosphate, GTP: guanosine 5’-triphosphate, NTP: nucleoside 5’-triphosphate with N = adenosine, guanosine, cytidine or uridine.

MSN play an important role in regulating bacterial growth ^[15,16]^. At the beginning of the exponential growth phase, ppGpp levels are rather low at ca. 40 µM^[15]^. During transition to stationary phase, a 20-fold increase occurs and it is generally assumed that the ppGpp levels increase in a linear fashion^[15]^. However, a mathematical description of bacterial growth during nutrient upshift predicted oscillations in different growth parameters, such as growth rate, ribosome allocation, ribosomal protein fraction and ppGpp content^[17,18]^. Even so, strong experimental data validating these predictions are missing for MSN concentrations during growth. Nevertheless, Teich et al. reported oscillating ppGpp levels during constant but limited addition of glucose to the medium^[19]^ and Friesen et. al.^[20]^ detected fluctuating ppGpp concentrations during nutrient upshift and downshift during the first 70 minutes of growth. Additionally, in *Bacillus subtilis* some fluctuations in ppGpp levels are detected during growth by live-imaging using an *in vivo* reporter construct^[21]^. In this assay, transcription of the firefly luciferase reporter is activated upon binding of (p)ppGpp to a regulatory sensor element positioned upstream of the luciferase gene. The emitted luminescence of the firefly luciferase provides a quantitative readout, with signal intensity correlating directly with intracellular MSN levels^[21]^. Unfortunately, the same study used only three time points for the validation of the MSN concentrations based on extraction/HPLC-determined levels. Another *in vivo* live-imaging assay was developed in *Escherichia coli*, but this study did not examine ppGpp levels during growth. The basis of this fluorescent read-out is the following: after binding of (p)ppGpp to an RNA-based biosensor, a Broccoli aptamer is associated with its cognate fluorophore. The increase in fluorescence can be detected and MSN levels can be measured^[22]^. However, both genetically encodable systems cannot distinguish between different (p)ppGpp species. The use of alternative aptamer classes may enable such discrimination in the future^[23]^. Extraction-based methods will still be required for validation and to exclude functional interferences caused by other related small molecules. Until now, the number of measurement points as well as the temporal resolution were also limiting in other extraction-based studies (e.g. Varik et al.^[15]^, Murray et. al.^[16]^), and therefore it remains to be analyzed whether ppGpp levels truly fluctuate during growth.

Despite intense research in the (p)ppGpp field^[24]^, the detection and quantification of (p)ppGpp is still a main issue because of levels close to the detection limit during unstressed growth^[7,15,25]^. Another problem is the difficulty in separation and assignment of different magic spot nucleotides^[23]^. Detection via mass spectrometry after UPLC separation was problematic for a long time, because high concentrations of nonvolatile phosphate buffers are often required for chromatography^[15,23,26]^. Nonetheless, methods using UPLC-MS are available^[27,28]^. One advantage of combining LC and MS is the possible differentiation of ppGpp and pppGpp by their masses, if separation of the two compounds is not achieved.

Whereas some methods are unable to distinguish between ppGpp and pppGpp, others are not sensitive enough to quantify or even detect the low abundance pppGpp^[15]^. This results in a potential underappreciation of pppGpp signaling, as the pentaphosphate is either not quantified or it is quantified together with ppGpp^[29]^. Nevertheless, in literature, there are HPLC^[28,30]^ and capillary electrophoresis (CE)^[31]^ methods discriminating between the tetra- and pentaphosphate with heavy internal standards like ^13^C_10_-(p)ppGpp, ^15^N_5_-(p)ppGpp or even a mix of ^13^C-(p)ppGpp and ^15^N-ppGpp^[28,30]^. Using these internal standards, MSN in principle do not have to be baseline separated (e.g. via HPLC in HILIC mode^[25]^ or CE^[31]^). In contrast, using mixed isotopic standards allows correction for the partial degradation of pppGpp to ppGpp that can occur during extraction, separation, or ionization^[30]^.

Another stress-responsive group of highly phosphorylated messenger molecules (alarmones) are dinucleoside polyphosphates (figure 1B). Structurally, this group is characterized by the connection of two nucleosides through an oligophosphate chain in between the 5’-OH groups. The general formula is Np_n_N (N = adenosine, guanosine, cytidine or uridine)^[32–34]^ with n usually varying between 2 to 6^[34,35,36]^, but even an Ap_7_A variant was reported in human platelets^[36]^. Np_n_Ns can be found in diverse organisms, with Ap_4_G, Ap_4_C, Ap_4_U and Ap_4_A being the most abundant variants in *E. coli*^[32,33]^. The exact signaling function of dinucleoside polyphosphates is not known; the molecules are synthesized by various enzymes like aminoacyl tRNA synthetases during starvation^[33]^ or by T4 RNA ligase^[37]^. Significant concentration jumps were found for example after oxidative stress^[38]^, suggesting that they indeed are part of the bacterial stress response. Basal levels of 0.2 – 1 µM Ap_4_A in *E*.*coli* during exponential growth were previously reported^[39,40]^, but time resolved changes in Ap_4_A levels were not published. Hydrolysis of Np_n_Ns is achieved by members of the NUDIX family of hydrolases or by ApaH, the main enzyme responsible for this process^[33,41]^. Interestingly, ApaH has recently been described to be inhibited by (p)ppGpp and thus a cross-talk between these alarmones seems likely^[42]^. Hence, an extraction protocol and quantitation method that allows monitoring of both Np_n_N and (p)ppGpp simultaneously would greatly facilitate studies into their functional connection.

Quantification of dinucleoside polyphosphates were performed for example by using bioluminescence^[45]^, 2-D thin layer chromatography^[40]^ or a chemiluminescence assay^[40]^. Recently, LC-MS/MS methods were published using ^13^C_20_-Ap_3/4_A internal standards for quantification of Ap_3/4_A^[39,46]^.

Given the need for accurate and precise quantitation methods in the alarmone field covering multiple analytes in a single analysis, we present a new method for quantifying magic spot nucleotides and dinucleoside polyphosphates in parallel. Using this method, we were able to detect fluctuations in the levels of magic spot nucleotides and dinucleoside polyphosphates as predicted mathematically^[17]^. These oscillations were shown before in *E. coli* under glucose-limiting conditions^[19,20]^, during nutrient upshift^[20]^, or in *B. subtilis* using live-imaging methods with luminescence read-out^[21]^. A confirmation with HPLC or CE coupled to MS was still missing. In mathematical descriptions by WU et al.^[47]^ calculations are based on radio TLC labeling during carbon limitation. Here, no fluctuations of ppGpp levels were detected. The data presented here could therefore further enhance such prediction models.

## Results

### Magic spot nucleotides and dinucleoside polyphosphate analysis by weak anion exchange (WAX) enrichment and capillary electrophoresis coupled to mass spectrometry (CE-MS)

Separation and quantitation of highly charged phosphorylated nucleotides is a challenging task, yet several different methods have been published^[23]^. The use of capillary electrophoresis (CE) enabled baseline separation (see figure 2B) of all phosphorylated purine analogues^[48]^. This approach, amended with a titanium dioxide (TiO_2_) extraction protocol, was already implemented in the quantification of ppGpp in plants^[31]^. However, the use of CE-MS for analysis after pre-spiking with heavy ^15^N_5_-(p)ppGpp references and weak anion exchange (WAX) extraction in bacteria has not yet been achieved and is one subject of this manuscript.

**Figure 2.**
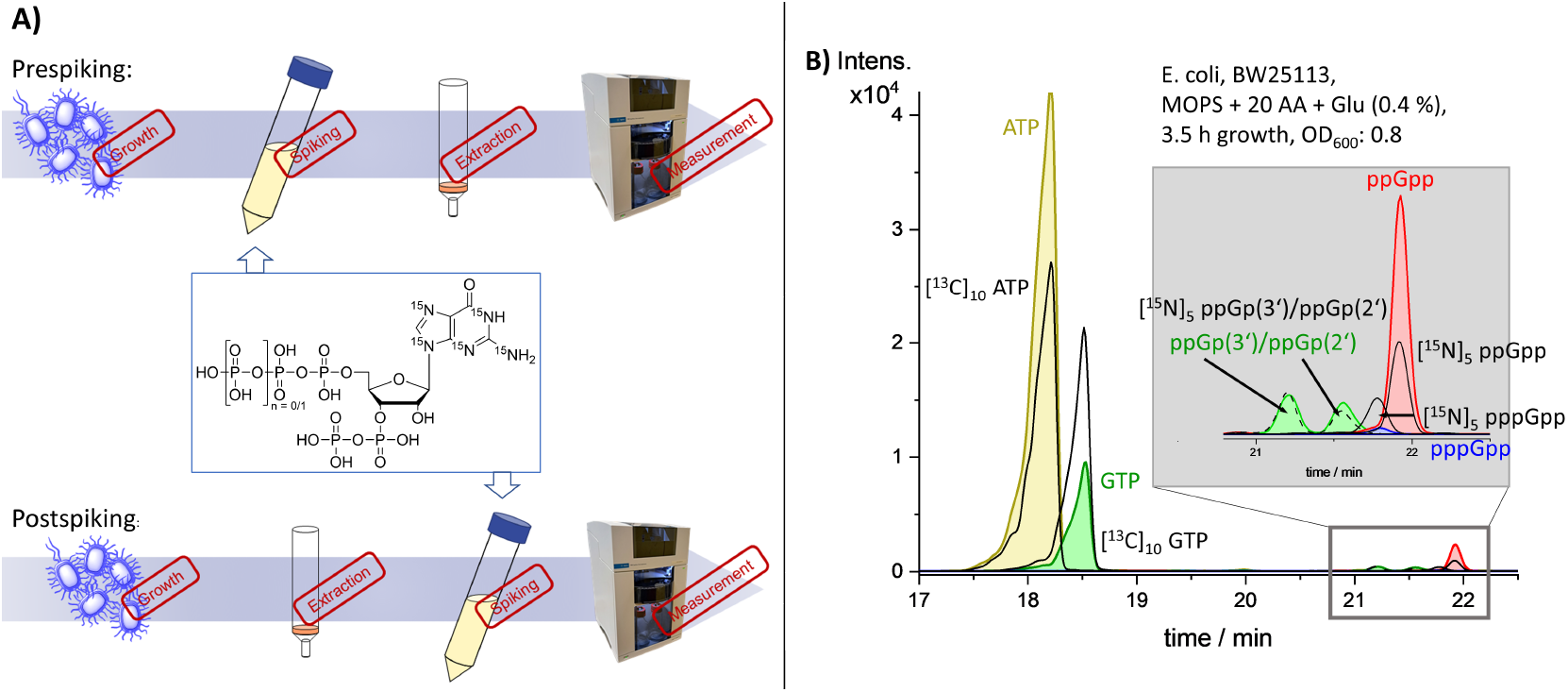
A) Workflow of pre- and post-spiking extractions. Unless specified otherwise, pre-spiking is used in all experiments. B) Electropherogram of ATP (yellow), GTP (green), ppGpp (red) and pppGpp (blue) with the corresponding heavy internal standards (black), analyzed by CE-MS. The peaks of ppGp(3’) and ppGp(2’) are also detectable after the dephosphorylation of ppGpp during incubation on ice (green peaks, box).

To test whether a baseline separation using CE can also be achieved from a bacterial matrix, the *E. coli* K 12 strain BW25113 was grown until an OD_600_ of 0.8 and 10 x 10^8^ bacterial cells were lysed and extracted, using a slightly modified version of the extraction protocol reported by Ihara et al.^[27]^ and Bartoli et al.^[28]^. The previously reported TiO_2_ extraction protocol was not applied because we observed a significant amount of 2’,3’-cyclophosphate formation and dephosphorylation of the pyrophosphate on the 3’-OH during TiO_2_ extraction compared to the WAX extraction. In the latter method, the negatively charged analytes interact with the positively charged amine resin and isolates anions from cationic and neutral metabolites. CE is highly sensitive to salt concentration, so the extraction of these analytes is critical for a successful implementation. The use of a dilution buffer with a higher pH (pH = 5.5 instead of pH = 4.5) compared to the published protocols^[27,28]^ facilitated a more stable pH of the bacterial extract, which was then applied to the cartridges (EVOLUTE® EXPRESS WAX, 100 mg/3 ml). This modification resulted in an improved peak shape in the subsequent analysis. The developed CE-ESI-MS/MS method^[31]^ enabled baseline separation of pppGpp, ppGpp, ATP and GTP also from a bacterial matrix with the automated WAX extraction protocol (figure 2B), underlining the separation efficiency of CE. Moreover, a separation of 2’-phosphate 5’-diphosphate (ppG(2’)p) and 3’-phosphate 5’-diphosphate (ppG(3’)p) is achieved. The dephosphorylation of the 3’-OH group and a phosphorylation of the 2’-OH of the ribose can occur during the incubation on ice during extraction via cyclophosphate formation and hydrolysis. The mechanism of the dephosphorylation to ppGp(3’) and ppGp(2’) is reported in Qiu et. al.^[31]^ and the respective peaks assigned to the analytes are shown in green in the electropherogram (box in figure 2B).

### Method validation with wild-type bacteria and a *relA* KO strain

Next, we tested the reproducibility of the protocol. Since MSN can readily decompose and lose one or more phosphates during incubation and extraction of MSN from the biological material^[30,31,48,49]^, pre-spiking of the lysed bacteria with stable isotope internal standards prior to extraction allowed us to compensate for this loss^[30,31]^. Such standards are now readily accessible by chemical synthesis^[31]^. The sample preparation and analysis workflow are shown in figure 2A. Prior to extraction, bacterial lysates were pre-spiked with a known amount of [^15^N]_5_-(p)ppGpp ^[31,48,50]^ and commercially available [^13^C]_10_-ATP and [^13^C]_10_-GTP followed by incubation on ice for 30 min. Six technical repeats at the end of exponential growth (OD_600_: 0.8) of wild-type *E. coli* grown in M63 medium showed stable levels of ppGpp (530 ± 90 pmol/10^8^ cells), pppGpp (106 ± 18 pmol/10^8^ cells), ATP (3080 ± 130 pmol/10^8^ cells), and GTP (1420 ± 150 pmol/10^8^ cells). The levels are reported in figure 3A. Amounts are indicated as pmol per 10^8^ cells rather than intracellular concentrations, because the cell size varies strongly during growth^[51]^. For comparison with literature, we estimated an average cell volume of 4 fL like described in Volkmer *et al*.^[52]^. Calculation of cellular ppGpp concentrations with these values gave ppGpp levels (∼ 1.3 mM) comparable to previous reports in the literature^[15]^ (0.8 mM) measured in MOPS medium.

**Figure 3.**
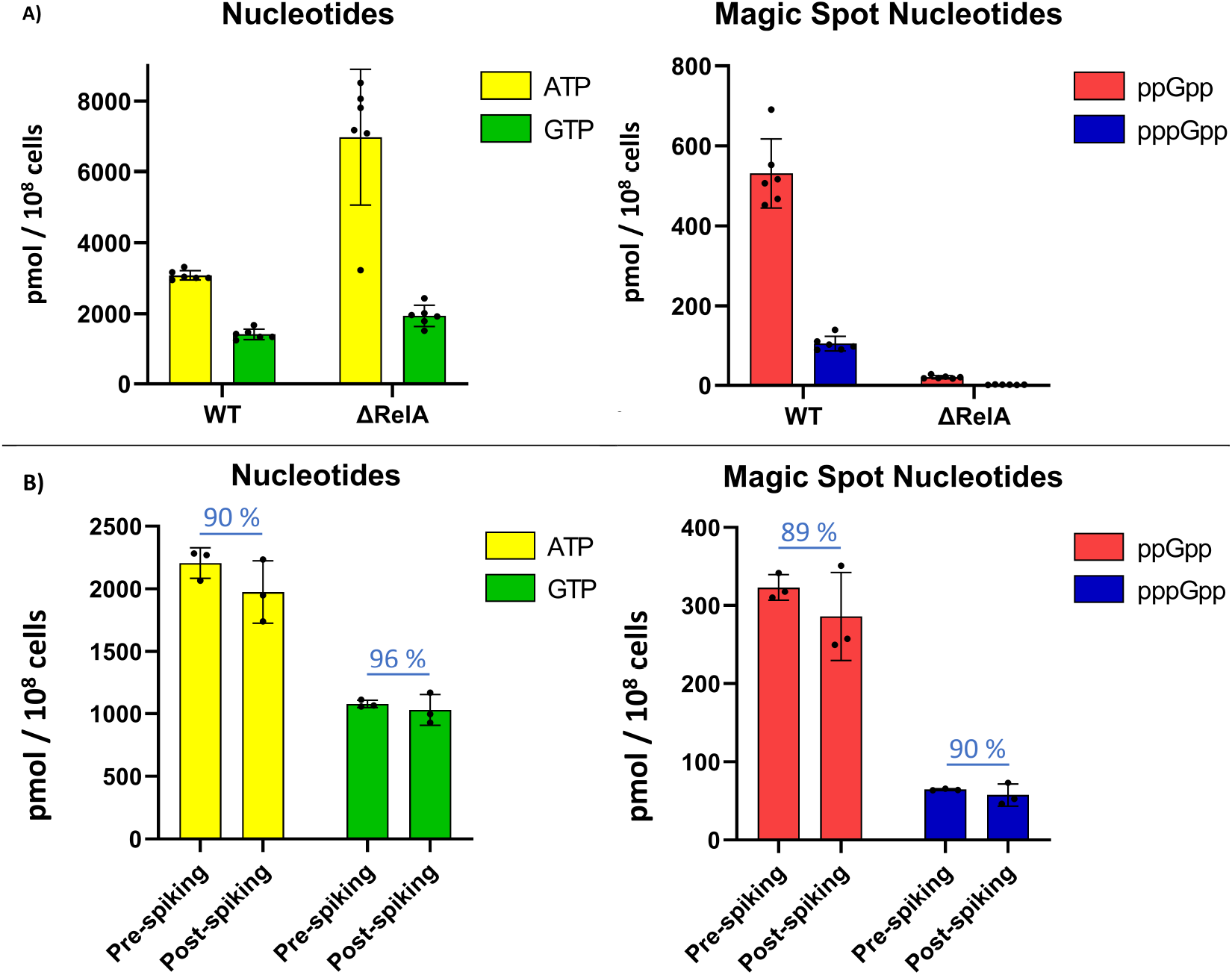
A) Levels of different nucleotides in validation experiments. B) Levels of different nucleotides in recovery experiments. Recoveries were calculated comparing levels of pre-spiking and post-spiking experiments. Error bars were calculated with GraphPadPrism using standard deviation. Nucleotides are depicted in the same colors as in Figure 2B).

As expected, levels of pppGpp were 5-fold lower in wild-type *E. coli* compared to ppGpp levels^[13]^ (figure 3A). In addition, both MSN were also detected and quantified in the *relA* KO strain, which lacks the main *E. coli* (p)ppGpp synthetase^[53]^. We detected a strong decrease of both MSN (ppGpp: 26-fold reduction, pppGpp: 67-fold reduction in the *relA* KO vs WT). Basal levels of ppGpp (20 pmol/10^8^ cells or 30 pmol/OD_460_) have been described previously for a *relA* KO strain^[13]^. Recalculation of the ppGpp levels measured at an OD_600_ of 0.8 showed a level of ppGpp in the *relA* KO of 26 pmol/OD_600_ comparable to previous literature reports. The remaining level of ppGpp is due to the activity of SpoT, a homolog of RelA^[5]^. SpoT is synthesizing (p)ppGpp after activation of the synthase domain, e.g. during carbon or fatty acid starvation^[29,54]^ although the main task of SpoT is likely the hydrolysis (p)ppGpp^[55]^.

### Determination of recovery in biological samples using heavy internal standards

It is important to know the extraction recovery of analytes from bacterial cell lysates and we tested two different work flows for comparison (see figure 2A). In all measurements, samples were spiked with defined amounts of heavy internal standards directly after cell lysis and thawing before extraction (pre-spiking). One can then track losses of the spiked nucleotides during extraction, because the loss of the internal heavy standards will be the same as the loss of the analyte. In another possible workflow, spiking of the samples was performed after extraction but before measurement (post-spiking). Here, losses during extractions are not quantifiable and have to be calculated by average values determined before. Determination of the recovery of analytes after the modified extraction procedure by Ihara et al.^[27]^ and Bartoli et al.^[28]^ were performed with six technical repeats split in two different sets. One set was pre-spiked and the other post-spiked and concentrations were determined and compared (figure 3B). Excellent recoveries of 89 ± 18 % for ppGpp, 90 ± 20 % for pppGpp, 90 ± 12 % for ATP and 96 ± 12 % for GTP were calculated, similar to those reported by Varik et al. using strong anion exchange (for ppGpp detection) or ion-paired reverse-phase chromatography (for ATP/GTP detection)^[15]^ and higher than the 40-70% recoveries reported for WAX extraction by Ihara^[27]^. As peaks of ppGp(3’) and ppGp(2’) can be detected in electropherograms (figure 2B), some of the losses of ppGpp (and pppGpp to pppGp(3’) and pppGp(2’), not shown) can be explained by the formation of a cyclophosphate diester involving the 2’- and 3’-OH of the ribose and a consecutive hydrolytic opening of this intermediate. However, the degradation during the incubation of (p)ppGpp in the beginning of the WAX extraction protocol is reduced compared to the degradation during a TiO_2_ extraction reported earlier^[31]^.

### Determination of magic spot levels during exponential and stationary phase in M63 medium

We monitored MSN and housekeeping nucleotide levels during bacterial growth in shorter time intervals (here: 0.5 h between sampling) than applied in other studies (e.g. Varik et al.^[15]^: after 2, 4, 6, 8, 10 h of growth). These shorter intervals were chosen to interrogate if MSN levels do indeed linearly increase in the exponential phase as often reported^[16,56]^ or oscillate like mathematically predicted^[17]^. We decided to take samples every 30 minutes, independent of the OD_600_. This approach was chosen to reduce cellular stress during sampling (e.g. temperature shifts), which could otherwise distort the measured alarmone levels. Several publications consistently report an increase in ppGpp levels in late exponential phase and a subsequent decrease in stationary phase^[15,16]^. In a first test experiment, bacteria were grown in M63 medium supplemented with all amino acids and glucose (0.4 %). As shown in figure 4A, levels in ppGpp were 2- to 6-fold higher than previously reported in another minimal medium (MOPS)^[15]^. The ppGpp levels started to increase in the middle of exponential growth. Moreover, fluctuations in ppGpp levels were noted in late exponential growth, which have been so far described only twice: during glucose limitation^[19]^ or during nutritional up- or downshift^[20]^. These fluctuations were explained by the non-synchronized cell states of bacteria^[19,57]^.

**Figure 4.**
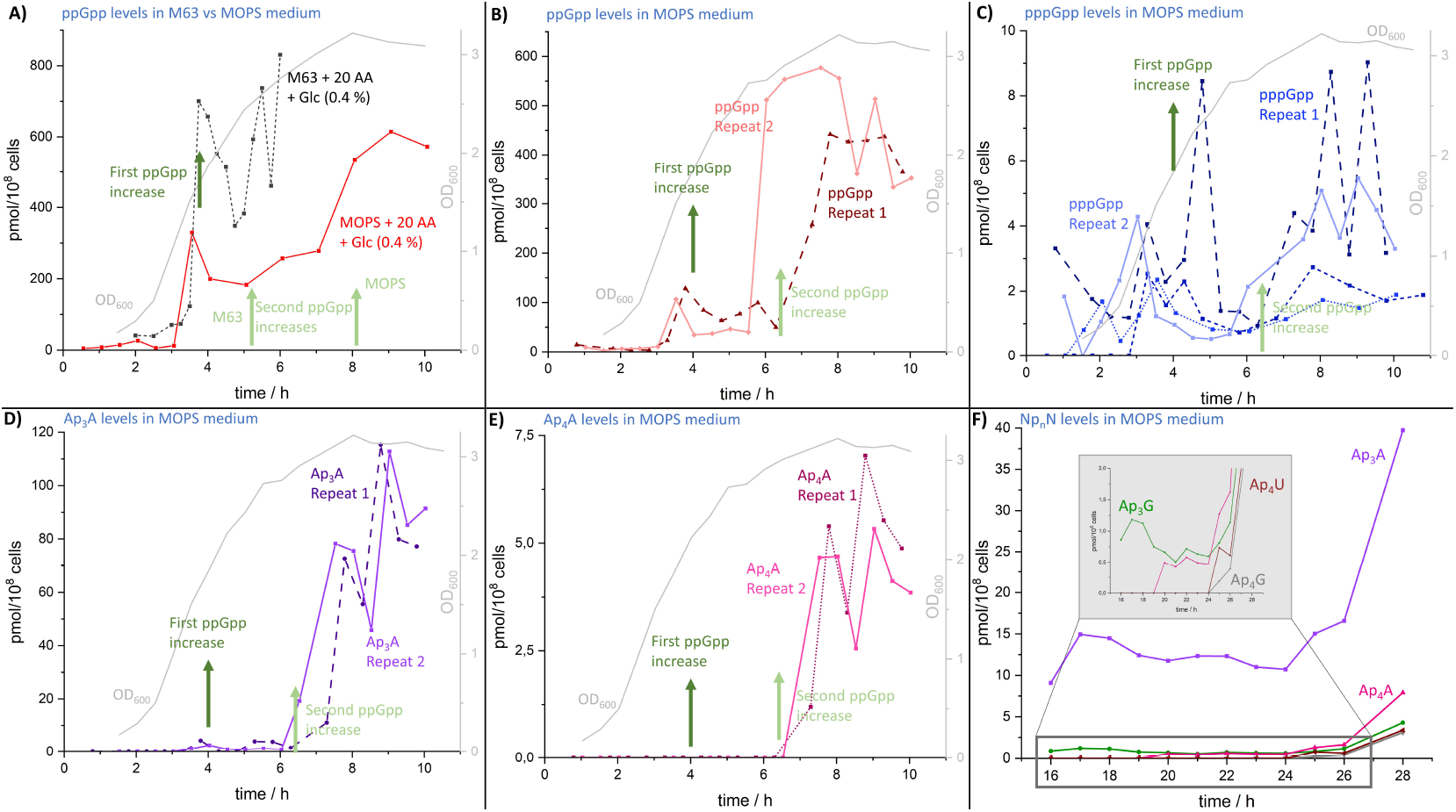
A) Levels of ppGpp during growth in two different growth media and the OD_600_ of one experiment of bacterial growth. In M63 (grey dotted line) higher levels of ppGpp are observable compared to ppGpp levels of bacteria grown in MOPS medium (red line). In both growth media, a first ppGpp peak (marked with a dark green arrow) is followed by a second larger increase (marked with light green arrow) in ppGpp levels. B) Levels of ppGpp (red line) of two different experiments in MOPS medium. In addition to a first increase in the middle of exponential growth (dark green arrow), a second increase (light green arrow) in stationary phase with higher levels compared to the first increase is observable. Later, oscillating levels of ppGpp can be detected. (Other repeats are shown in figure S1). C) Fluctuating levels of pppGpp. (Other repeats are shown in figure S4.) D) Ap_3_A levels during exponential growth of two different repeats. An increase of Ap_3_A levels comparable to ppGpp is observable. The observed oscillations of ppGpp are also visible in Ap_3_A levels. E) Ap_4_A levels of two different experiments during growth. After a first increase, Ap_4_A levels fluctuate. F) Np_n_N levels during 16 to 28 hours of growth. *E. coli* grown in M63 medium supplemented with glucose (0.4 %) and 18 amino acids without supplementation of methionine and cysteine. Shown are Ap_3_A in purple, Ap_3_G in green, Ap_4_A in pink, Ap_4_G in grey, Ap_4_U in brown. B)-E): Increases of ppGpp are marked with a dark green (first increase) and a light green arrow (second increase). Repeat 1 is represented by a dotted line and in dark colors, repeat 2 by a line and light colors. In grey, OD_600_ of one experiment is shown for comparison of bacterial growth. For better comparison, the time scale was adjusted so that the turning point in OD_600_ measurements occurred at the same time.

### Determination of magic spot levels during exponential and stationary phase in MOPS

Varik et al. used a supplemented MOPS medium^[15]^ and not the M63 medium reported here. M63 medium contains more phosphate and ammonium, MOPS medium is instead buffered with morpholinopropane^[58]^. The question was, whether the observed ppGpp oscillations were a specific response to the medium or whether they also would occur in MOPS medium. Therefore, nucleotide levels were redetermined for bacteria grown in MOPS medium. ppGpp levels were 2-fold lower during exponential growth compared to M63 medium and closer to levels published earlier^[15,25]^, but again increased during mid-exponential phase. Importantly, the oscillations were also reproducibly observed in supplemented MOPS medium, although the differences in the absolute levels were not as pronounced as in the supplemented M63 medium (see figures 4A, 4B and S1). In conclusion, the data presented here are an experimental validation of the mathematically predicted oscillating ppGpp levels^[17]^. In these predictions, a feedback system comprising a sensor and an effector was proposed as the mechanism driving the oscillations. Another approach showed that allocating resources to gene expression and precursor production - known as the *precursor-only strategy* - can also generate oscillations in ribosomal protein synthesis and ppGpp levels^[18]^.

Differences in absolute ppGpp concentrations can potentially be explained with mutation rates of *E. coli*^[59]^. The used K-12 strain often acquires spontaneous mutations and mutant alleles of *relA* and *spoT* can influence the (p)ppGpp levels^[10]^. Importantly, the turnover rates of ppGpp and pppGpp are on the order of seconds to minutes^[15,60]^.

### Levels of pppGpp during growth

The oscillations observed in ppGpp levels were also reflected in those of pppGpp (figure 4C). However, the characteristic pattern - an initial rise followed by fluctuating levels - was less pronounced for pppGpp. The concentration of pppGpp increased rapidly during early exponential growth, then declined to minimal levels in late exponential phase, before rising again upon entry into stationary phase (figure 4C). This effect may result from the generally lower cellular abundance of pppGpp and its faster turnover rate. Such oscillations are of regulatory interest, since ppGpp and pppGpp can affect different cellular targets at different concentrations^[7,29,61]^.

### Detection of Ap_3_A and Ap_4_A

Ap_3_A and Ap_4_A are members of the dinucleoside polyphosphate family of alarmones and are among the most thoroughly investigated within this group^[62]^. From the same extracts obtained as described before, we were also able to detect and quantify Ap_3_A (recovery of Ap_3_A standards in aqueous solution: 100 µM: 91.2 ± 1.8 %, 10 µM: 99 ± 6 %) and Ap_4_A (recovery of Ap_4_A standards in aqueous solution: 100 µM: 99 ± 11 %, 10 µM: 73.9 ± 1.6 %, method: see SI, page 15).

Peak assignment of Ap_3_A (figure S17) was performed using a heavy ^18^O internal standard synthesized in our lab according to a published procedure^[63]^ (high resolution mass spectrometry (HRMS) ([M-H]^-^, found: 755.0734, calculated: 755.0747, see table S1) and fragmentation pattern (see methods, table 2). Ap_4_A peak assignment was achieved with HRMS ([M-2H]^2-^, found: 417.0169, calculated: 417.0169, see table S1) and a characteristic fragmentation pattern (see methods, table 2). For Ap_4_A, a heavy internal standard^[46]^ was not available in our laboratory and therefore quantification of Ap_3_A and Ap_4_A levels were ultimately determined using the signal of the heavy ppGpp internal standard, as the singly ^18^O-labeled Ap_3_A was ambiguous. Nonetheless, using the heavy Ap_3_A standard, the limit of detection (LOD) for Ap_3_A was determined to be 34 nM and the limit of quantification (LOQ) to be 0.1 µM (method see SI, page 13). The LOD and LOQ of Ap_4_A were calculated to 2.0 ± 0.3 µM and 6.8 ± 1.2 µM, respectively. No heavy Ap_4_A standard was available, so a linear regression of different concentrations vs abundances of detectable Ap_4_A peaks were used (method see SI, figures S15 and S16).

### Ap_3_A and Ap_4_A accumulation during in *E. coli* growth

The first increase of Ap_3_A to detectable levels occurred during growth at an OD_600_ > 1.5 (ca. after 3.5 hours). In all measurements, this first Ap_3_A spike was detected during mid-to late exponential phase (approx. 6 h, see figure 4D). During stationary phase, levels are increasing and show oscillations in the range of 50 -100 pmol per 10^8^ cells. Interestingly, these changes of the Ap_3_A levels during growth coincided with changes in ppGpp levels (marked with green arrows in figure 4D), which could indicate that their synthesis is coordinated.

Comparison with literature is not possible because Ap_3_A concentrations were so far only reported for cells grown in MOPS medium to early exponential phase (0.1 ≤ OD_600_ ≤ 0.5)^[64]^. During early exponential phase, Ap_3_A concentrations are below our LOD, which likely reflects that we used about 4-8 times less bacteria than in the study by Plateau et. al.^[64]^. We calculated the intracellular concentrations in order to compare the Ap_3_A levels measured at higher OD_600_ values with the reported ones during early growth. As Ap_3_A levels were not stable (figure 4D), average levels of Ap_3_A after the first detection (OD_600_: 1.5) until the end of exponential growth (OD_600_: 2.8) were used for comparison. With an average Ap_3_A level of 1.78 pmol/10^8^ cells and an estimated cell volume of 4 fL^[52]^, an intracellular Ap_3_A concentration of 4.5 µM was calculated. This concentration is 10-fold higher than the Ap_3_A cellular concentration previously reported for lower OD_600_ values in the same medium (0.5 µM)^[64]^ and further underscores the significant increase in Ap_3_A levels in late exponential growth phase.

The first occurrence of an Ap_4_A concentration spike was during transition of late exponential to stationary phase (OD_600_: 3) and appeared coordinated with an Ap_3_A spike (figure 4E). In the stationary phase, comparable Ap_4_A fluctuations as reported for ppGpp and Ap_3_A were detectable. These coordinated spikes make it unlikely that diadenosine polyphosphates are just mere unavoidable byproducts of other biochemical transformations.

Reported Ap_4_A concentrations in different *E. coli* strains (MG1655^[39]^ or AB1157^[40]^) and in minimal or complex medium (M9^[39]^or LB^[40]^) during early exponential growth (OD_600_: 0.3^[39]^ and OD_600_: 0.3^[40]^) range from 0.2 to 1 µM^[39,40]^. *E. coli* grown in MOPS medium, however, showed increased Ap_4_A levels of 2.4 µM (OD_600_: 0.5)^[32]^ and 3.2 (OD_600_: 0.3 - 0.5)^[45]^, respectively. Corresponding values for stationary phase have not yet been determined. The Ap_4_A levels in *E. coli* BW25113 were below LOD during early and mid-exponential growth in our experiments, but Ap_4_A levels found in the stationary phase (ca. 12 µM, calculation parameters see above) are 4- to 60-fold higher than the levels reported in literature during exponential phase^[32,39,40,45]^.

### Coordinated oscillations of ppGpp, Ap_3_A and Ap_4_A levels in *E. coli*

During the transition from exponential growth to stationary phase, we consistently detected a second and higher spike of Ap_3_A levels (figure 4D), comparable to the increase of the magic spot nucleotides. An identical pattern was found for Ap_4_A (figure 4 F). As for the magic spot nucleotides, these Ap_3_A and Ap_4_A oscillations likely indicate a fast turnover rate. The synchronized behavior of the levels of these alarmones suggests similar regulation and potentially synergistic functions. The earliest increase in ppGpp levels could indicate that during the stress response of bacteria the synthesis of dinucleoside polyphosphates is stimulated, or degradation is reduced, for example by inhibition of the hydrolase ApaH by (p)ppGpp^[42]^. Spikes of Ap_3_A might then lead to its use as a noncanonical initiator nucleotide as reported before^[41,65]^ and might initialize the increase of capped and stabilized^[65]^ RNA during the stationary phase of *E. coli*^[41]^.

### Np_n_N levels increase during late exponential phase

Magic spot nucleotides as stress signals are more abundant in stationary phase, where stress factors become more prevalent. We were also interested in levels of dinucleoside polyphosphates after longer growth times and analyzed their cellular abundance between 16 to 40 hours (figure 4F). To obtain higher absolute concentrations of Np_n_Ns for ease of detection, experiments were performed in M63 medium under methionine and cysteine starvation, but with the supplementation of all other amino acids and glucose. After 16 hours of growth in M63 minimal medium without the supplementation of the above-mentioned amino acids, Ap_3_A was the most abundant dinucleoside polyphosphate with stable levels until 26 hours of growth (figure 4F). Ap_3_G levels were ca. 10-fold lower and stable. During the analyzed time period, three other dinucleoside polyphosphates were detected: Ap_4_A, Ap_4_G and Ap_4_U. After 26 hours and up to 40 hours of growth, an increase of all dinucleoside polyphosphates (Ap_3_A, Ap_3_G, Ap_4_A, Ap_4_G and Ap_4_U) was observed. Moreover, during this time period, we observed a 12-fold decline of ppGpp levels (see figure S2). This could be an indication of a changing control mechanism of the (starvation) stress response of *E. coli*: the initial stress response is organized by MSN, but during subsequent starvation stress, dinucleoside polyphosphates become more abundant and important.

All Np_n_N levels after 16, 26 and 40 hours of growth are summarized in table 1. Moreover, time-points and levels of their first detection and the increases between 16 and 40 hours (Ap_3_A, Ap_3_G) or after first detection (Ap_4_A, Ap_4_G and Ap_4_U) of growth are shown. The identities of the dinucleoside polyphosphates were assigned by high resolution mass spectrometry using calculated masses (see table S1).

**Table 1:** Levels of Np_n_N after 16, 26 and 40 hours of growth in M63 medium supplemented with 18 amino acids (-methionine, - cysteine) and glucose. All levels are indicated in pmol/10^8^ cells.

|  | Ap<sub>3</sub>A | Ap<sub>3</sub>G | Ap<sub>4</sub>A | Ap<sub>4</sub>G | Ap<sub>4</sub>U |
| --- | --- | --- | --- | --- | --- |
| **Np<sub>n</sub>N level first detection** | 4.28 | 0.85 | 0.49 | 0.19 | 0.74 |
| **Time of first detection** | 3.5 | 16 | 20 | 25 | 25 |
| **Np<sub>n</sub>N level after 16 h** | 9.09 | 0.85 | 0.00 | 0.00 | 0.00 |
| **Np<sub>n</sub>N level after 26 h** | 12.42 | 1.14 | 1.62 | 0.39 | 0.61 |
| **Np<sub>n</sub>N level after 40 h** | 340.61 | 340.43 | 52.75 | 49.48 | 34.58 |
| **fold increase\*** | 37 | 395 | 107 | 258 | 47 |

\* Increases calculated between first detection (Ap<sub>3</sub>A, Ap<sub>4</sub>A, Ap<sub>4</sub>G and Ap<sub>4</sub>U) or after 16 h (Ap<sub>3</sub>G) and 40 h of growth.

**Table 2:**
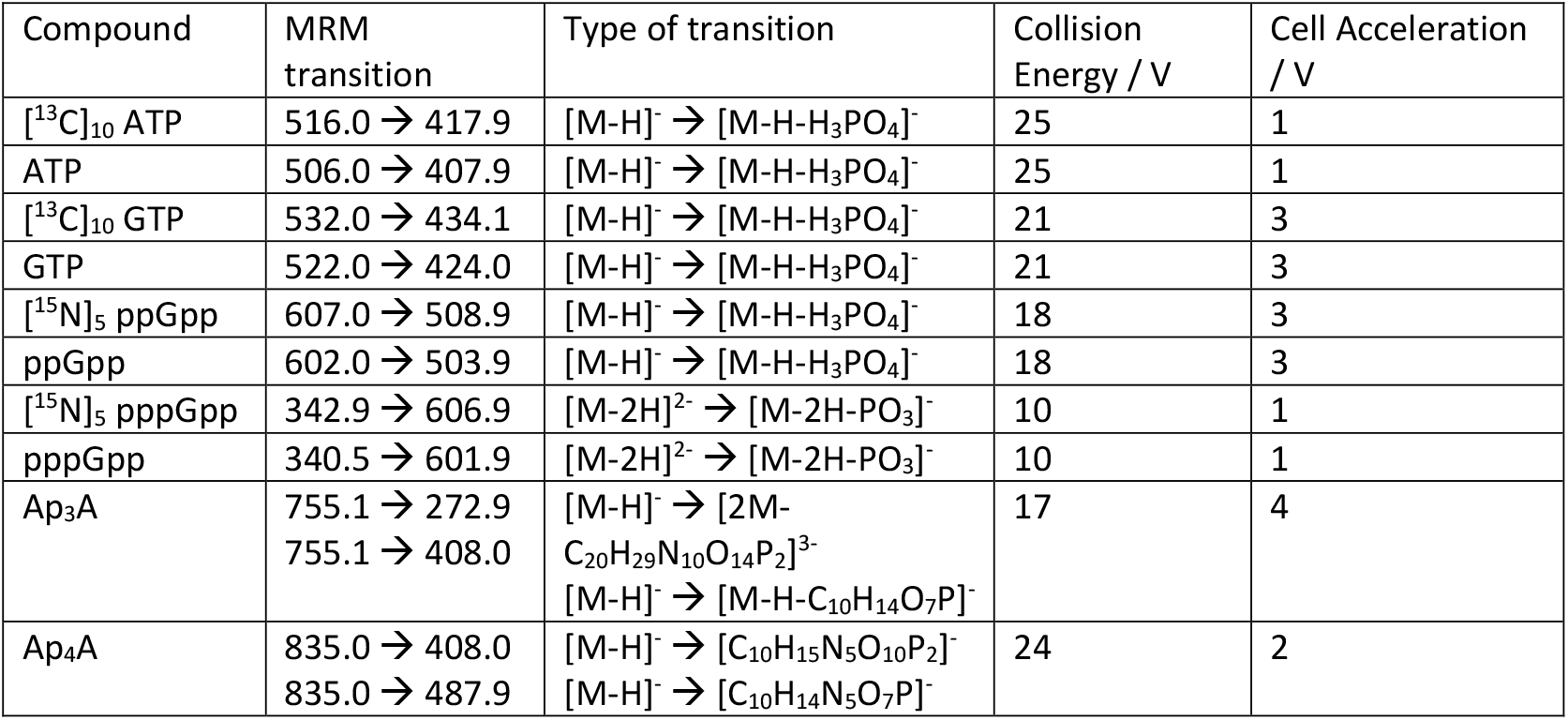
MS/MS transitions, collision energies and cell acceleration of compounds measured on Agilent G6495C Agilent QQQ.

| Compound | MRM transition | Type of transition | Collision Energy / V | Cell Acceleration / V |
| --- | --- | --- | --- | --- |
| [<sup>13</sup>C]<sub>10</sub> ATP | 516.0 → 417.9 | [M-H]<sup>-</sup> → [M-H-H<sub>3</sub>PO<sub>4</sub>]<sup>-</sup> | 25 | 1 |
| ATP | 506.0 → 407.9 | [M-H]<sup>-</sup> → [M-H-H<sub>3</sub>PO<sub>4</sub>]<sup>-</sup> | 25 | 1 |
| [<sup>13</sup>C]<sub>10</sub> GTP | 532.0 → 434.1 | [M-H]<sup>-</sup> → [M-H-H<sub>3</sub>PO<sub>4</sub>]<sup>-</sup> | 21 | 3 |
| GTP | 522.0 → 424.0 | [M-H]<sup>-</sup> → [M-H-H<sub>3</sub>PO<sub>4</sub>]<sup>-</sup> | 21 | 3 |
| [<sup>15</sup>N]<sub>5</sub> ppGpp | 607.0 → 508.9 | [M-H]<sup>-</sup> → [M-H-H<sub>3</sub>PO<sub>4</sub>]<sup>-</sup> | 18 | 3 |
| ppGpp | 602.0 → 503.9 | [M-H]<sup>-</sup> → [M-H-H<sub>3</sub>PO<sub>4</sub>]<sup>-</sup> | 18 | 3 |
| [<sup>15</sup>N]<sub>5</sub> pppGpp | 342.9 → 606.9 | [M-2H]<sup>2-</sup> → [M-2H-PO<sub>3</sub>]<sup>-</sup> | 10 | 1 |
| pppGpp | 340.5 → 601.9 | [M-2H]<sup>2-</sup> → [M-2H-PO<sub>3</sub>]<sup>-</sup> | 10 | 1 |
| Ap<sub>3</sub>A | 755.1 → 272.9<br>755.1 → 408.0 | [M-H]<sup>-</sup> → [2M-C<sub>20</sub>H<sub>29</sub>N<sub>10</sub>O<sub>14</sub>P<sub>2</sub>]<sup>3-</sup><br>[M-H]<sup>-</sup> → [M-H-C<sub>10</sub>H<sub>14</sub>O<sub>7</sub>P]<sup>-</sup> | 17 | 4 |
| Ap<sub>4</sub>A | 835.0 → 408.0<br>835.0 → 487.9 | [M-H]<sup>-</sup> → [C<sub>10</sub>H<sub>15</sub>N<sub>5</sub>O<sub>10</sub>P<sub>2</sub>]<sup>-</sup><br>[M-H]<sup>-</sup> → [C<sub>10</sub>H<sub>14</sub>N<sub>5</sub>O<sub>7</sub>P]<sup>-</sup> | 24 | 2 |

### Np_n_N levels without supplementation of two amino acids in the growth medium

As dinucleoside polyphosphate levels were strongly increasing after 26 hours of growth, we decided to also determine their levels after 36 - 56 hours of growth (figure 5A). Again, we used M63 medium supplemented with glucose and 18 amino acids, i. e. without methionine and cysteine.

**Figure 5.**
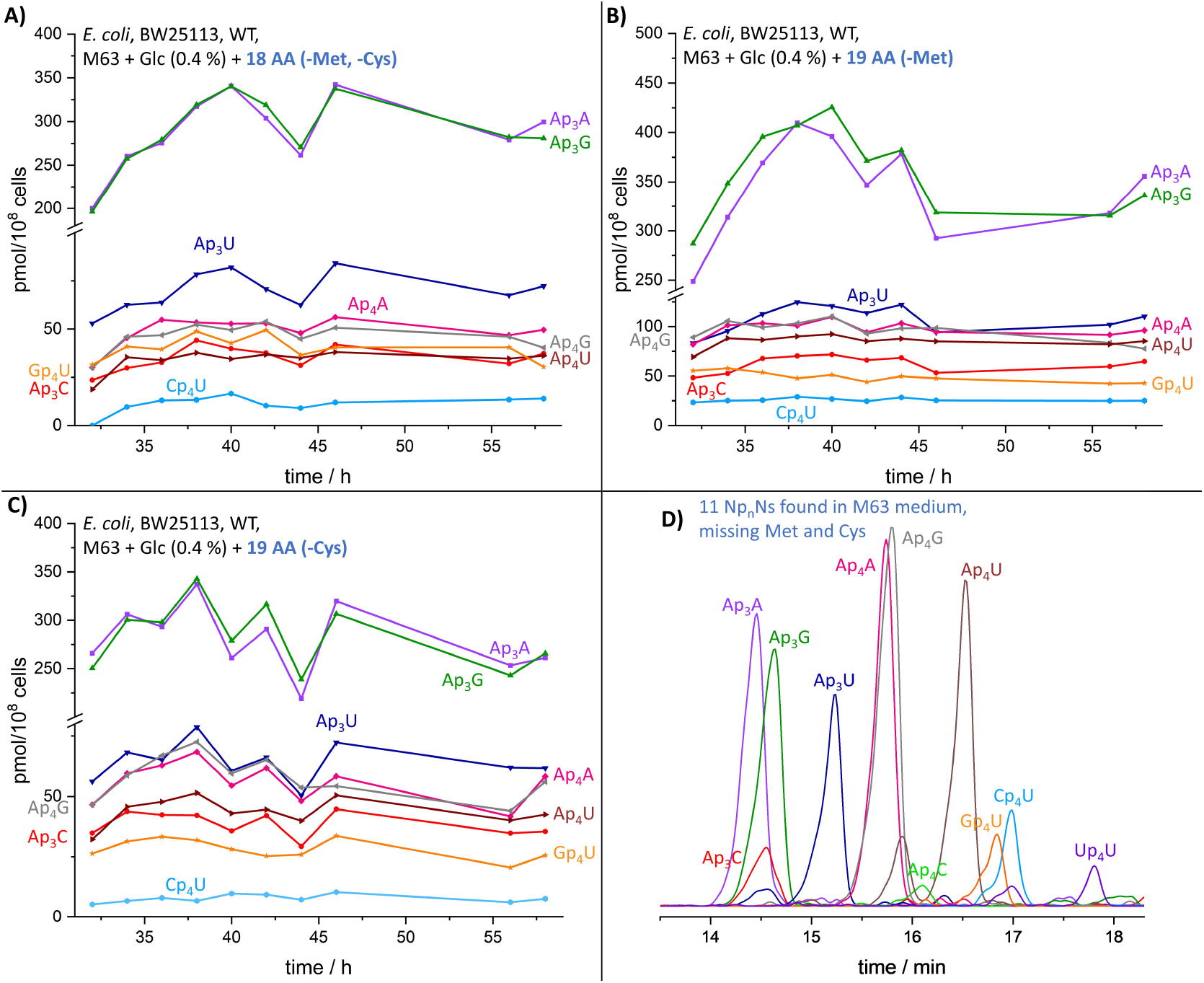
Np_n_N levels after 32 to 58 hours of growth with varying amino acid supplementation. All media contained M63 and glucose (0.4 %). Shown are Ap_3_A in purple, Ap_3_C in red, Ap_3_G in green, Ap_3_U in blue, Ap_4_A in pink, Ap_4_G in grey, Ap_4_U in brown, Gp_4_U in orange. A) Supplementation of 18 amino acids excluding methionine and cysteine. B) Supplementation of 19 amino acids without methionine. C) Supplementation of 19 amino acids excluding cysteine. D) Electropherogram of all detected Np_n_Ns after 24 hours of growth in M63 medium without supplementation of methionine and cysteine, but with all other amino acids and glucose (0.4 %). The following dinucleoside polyphosphates were detected in this sample: Ap_3_A (light purple), Ap_3_C (red), Ap_3_G (dark green), Ap_3_U (dark blue), Ap_4_A (pink), Ap_4_C (light green), Ap_4_G (grey), Ap_4_U (brown), Cp_4_U (light blue), Gp_4_U (orange) and Up_4_U (dark purple); exact recoveries are only available for Ap_3_A and Ap_4_A.

Eight dinucleoside polyphosphates were detected according to their high-resolution masses at high levels (figure 6, grey). Ap_3_A and Ap_3_G were most abundant, whereas Ap_4_A, Ap_4_G, Ap_4_U, Gp_4_U, Ap_3_C, and Ap_3_U were detected at lower concentrations. The levels of these alarmones consistently increased up to 46 hours of growth. The levels of each Np_n_N were much higher (grey arrows in figure 6) in cells grown under amino acid starvation compared to cells grown in M63 medium containing all amino acids (figure 6, blue). Thus, amino acid starvation induces dinucleoside polyphosphate synthesis. As dinucleoside polyphosphates increased, ADP and ATP levels, serving as precursors of adenosine nucleoside polyphosphates, declined (ATP: ca. 58-fold, ADP: ca. 5-fold). These nucleotides reached their lowest point after 58 hours of growth (see figure S9 and S11). Levels of Np_n_Ns after 36 to 58 hours of growth with the supplementation of all 20 amino acids are shown in the supporting information (figure S5).

**Figure 6.**
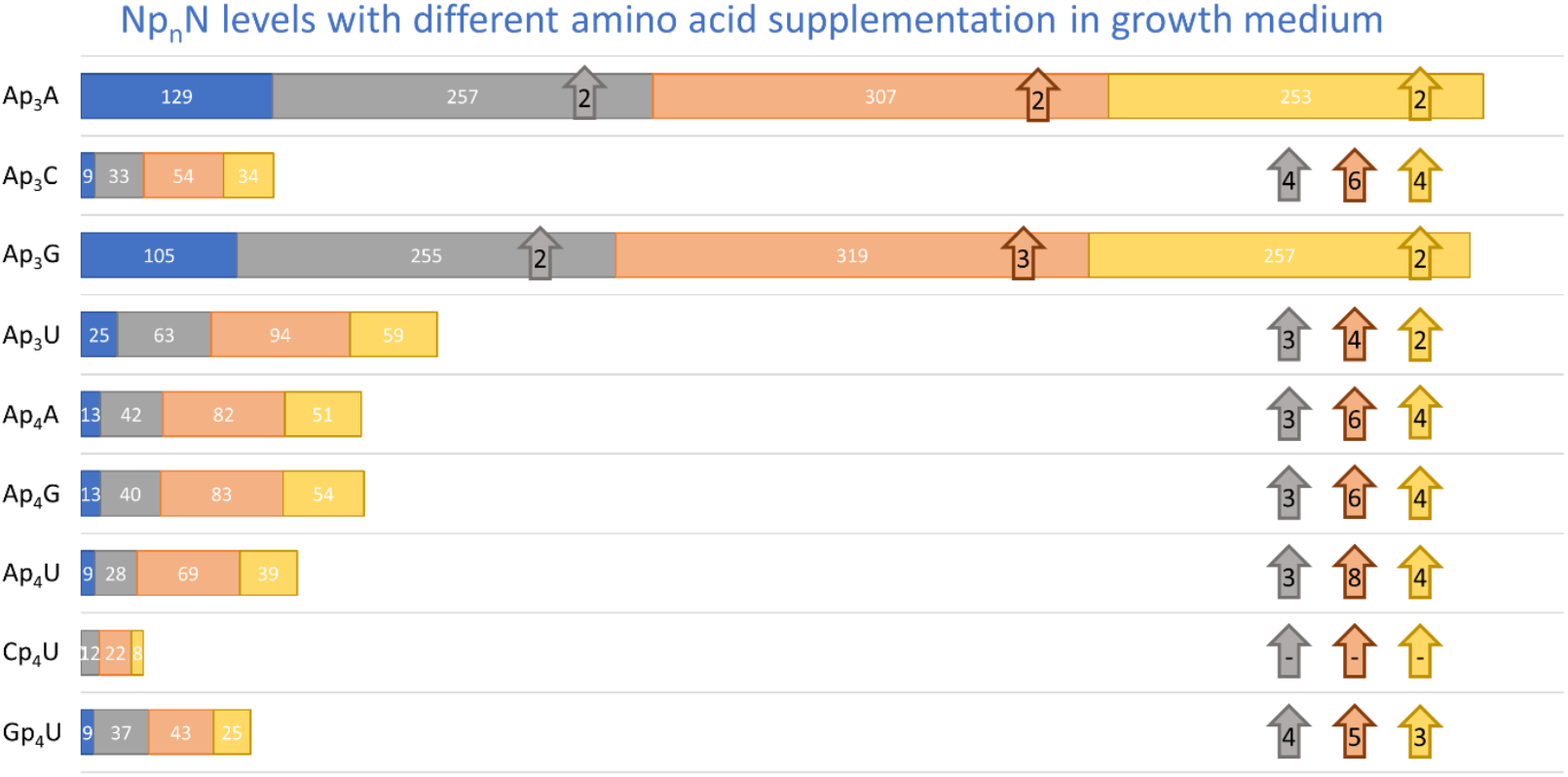
Np_n_N levels (in pmol/10^8^ cells) of *E. coli* grown in medium supplemented with 20 amino acids (blue), 19 amino acids (-methionine: orange; or -cysteine: yellow) and 18 amino acids (-methionine, - cysteine: grey). Arrows indicate the fold changes in Np_n_N levels between the growth in different media conditions compared to growth in medium supplemented with 20 amino acids (arrow colors like bars).

### Np_n_N levels in cells grown in the absence of a single amino acid

Both methionine and cysteine are important amino acids as methionine serves as initiating amino acid during translation and cysteine is required for disulfide-bridge formation and thus folding. We wanted to analyze, whether the observed increase in the Np_n_N levels reflect such specific functions or whether their individual absence causes the same effect. The dinucleoside polyphosphate levels were determined during a time period of 32 - 58 hours of growth in the M63 medium lacking just methionine (figure 5B). Interestingly, higher levels of all quantified dinucleoside polyphosphates were detected during methionine starvation compared to all other conditions (figure 6, orange). The increases in methionine starvation compared to growth medium supplementation with all 20 amino acids are quantified with orange arrows in figure 6. This increase in the synthesis of dinucleoside polyphosphates might be an indication of the special importance of methionine. Interestingly, ppGpp levels remain stable in (very) late stationary phase with all tested supplementations (figure S3), and so a decoupling of these alarmones is apparent.

Another unique amino acid is cysteine, which can form disulfide bonds, important for the stability and folding of enzymes^[66]^. Bacteria grown in medium without cysteine supplementation showed lower (figure 6C) levels of Np_n_Ns compared to bacteria without methionine supplementation, and comparable levels to the supplementation of 18 amino acids missing methionine and cysteine. The lack of cysteine in the growth medium seems to have less consequences compared to the lack of methionine. In figure 6, highest levels of Np_n_N without cysteine supplementation are shown in yellow. Increases of Np_n_Ns with this growth medium supplementation compared to the supplementation with 20 amino acids are indicated with yellow arrows. Interestingly, the omission of cysteine appears to slightly reduce the effects of methionine deficiency in the medium. The levels of dinucleoside polyphosphates (figure 6) are about the same as in bacteria grown without cysteine and methionine supplementation (figure 5A/C, figure 6). The increases of dinucleoside polyphosphates without cysteine (figure 6) and of 18 amino acids (2 - 4-fold, compared to 20 amino acid supplementation, respectively) are in the same range. In figure 6, the highest levels of all detected dinucleoside polyphosphates are shown, as well as the increase of Np_n_Ns during methionine starvation compared to the medium supplementation with all amino acids (yellow arrows).

In all of the tested conditions, levels of Ap_3_A and Ap_3_G were higher compared to the other dinucleoside polyphosphates found. As BENONI ET AL.^[65]^ previously reported, Ap_3_A/G attachment to the 5’-end of bacterial mRNAs promote both, transcription initiation and mRNA stability. Our results provide further indication of the use of Ap_3_A/G as noncanonical initiator nucleotides in (very) late stationary phase, based on their relative abundance.

### 11 Np_n_Ns found in amino acid starved *E. coli*

The levels of other Np_n_Ns were below the detection limit until 30 hours of growth. To gain further insights into other possible dinucleoside polyphosphates in *E. coli*, 1.4-fold more material (up to 40 × 10^8^ cells) was extracted. The extracted bacteria (BW25113) were grown in M63 medium supplemented with glucose (0.4%) and 18 amino acids (lacking methionine and cysteine). In this sample, two additional dinucleoside polyphosphates, Ap_4_C and Up_4_U were detected (see figure 5D). The newly formed dinucleoside polyphosphates were previously detected after temperature shift or after exposure of *E. coli* to cadmium and also during exponential growth measured by bioluminescent analysis^[32,64]^. The relative concentrations of all detected Np_n_Ns are summarized in figure S6.

Probably, dinucleoside polyphosphates can be detected during (very) late stationary phase in the same medium in a larger amount because the metabolism of *E. coli* changes during starvation. The increase of these metabolites might initialize the increase of capped RNA^[65]^ during the stationary phase of *E. coli*. Moreover, with the strong increase of Np_n_Ns in late stationary phase, ppGpp levels decrease. This could be an indication of Ap_4_A as another growth arrest signal. In fact, these compounds, especially Ap_3_A and Ap_4_A, were defined as damage metabolites occurring in high concentrations in different critically damaged cells^[34]^.

## Conclusion

Magic spot nucleotides and dinucleoside polyphosphates are known since 1969^[3]^ and 1966^[67]^, respectively. Nevertheless, the signaling of Np_n_Ns and the interaction between these messenger families are still not fully understood and are only rarely explored. Little is known about changes in Np_n_N levels during growth. Furthermore, only a few HPLC methods^[27,28,30,39,46]^ for both families and two methods for *in vivo* quantification^[21,22]^ of MSN are available. These methods have not been developed to determine MSN and Np_n_N levels in parallel. Herein, we developed an extraction and analysis method for the parallel quantification of diverse alarmones: MSN and Np_n_N. Applying this method, fluctuations in levels of both groups and sharp increases in all Np_n_N levels were detected that show initial coupling and then a decoupling in very late stationary phase.

Our CE-MS method with heavy internal standards shows stable levels of (p)ppGpp levels in technical repeats of wild type *E. coli* and a knockout mutant of RelA. Using a pre- and post-spiking method with those standards, a high recovery (around 90%) of analytes from extractions was confirmed^[15]^. Application of the new method to determine the levels of MSN during growth in M63 medium revealed significantly higher and oscillating levels of (p)ppGpp than reported before in MOPS medium^[15,16]^. With the change of M63 to MOPS medium levels of (p)ppGpp decreased to comparable levels like reported before^[15]^, but the oscillations were still observable. What could be the reasons for such oscillations? One assumption is that after the onset of starvation, the stringent response starts and RelA is synthesizing MSNs. With the increase of ppGpp, RelA is further activated^[68]^. A too substantial increase of MSN is toxic^[7]^ and is inhibited by the counterpart of RelA: SpoT^[13]^. Mathematical calculations indicate, that SpoT is not only hydrolyzing MSNs in a “simple term”^[17]^, but is modulated. A too massive overshoot is therefore prevented, resulting in oscillations. Another possibility is related to the switch between inactive and active state of ribosomes: Giordano et al.^[18]^ were able to show that an optimized growth rate is obtained in response to nutrient supply by switching^[17]^. As ppGpp levels are directly linked to active and inactive ribosomes^[18]^ these ribosome oscillations are potentially transferred to ppGpp concentrations. Finally, the fluctuations could be necessary as a control, whether the bacterial growth or dormancy is still in accordance to the nutrient supply. With decreasing MSN levels, bacterial growth can restart. If the lack of nutrients still exists, the synthesis of (p)ppGpp can restart. As a result, optimal tuning of nutrient supply vs growth is achieved by the help of oscillations.

Our experiments with short sampling intervals strongly support the occurrence of (p)ppGpp oscillations during exponential growth - a subtle feature that is usually overlooked but may have profound implications, as discussed above. These oscillations may be communicated to other alarmones, such as Np_n_N, e.g. through inhibition of ApaH by (p)ppGpp^[42]^. Apparently, this connection between concentrations of alarmones is lost during very late stages of growth and up to 11 Np_n_N appear as stress metabolites. A key amino acid for Np_n_N synthesis might be methionine, as the dinucleoside polyphosphate levels were the highest during methionine starvation in three tested amino acid supplementations. The strongest increase (roughly 400-fold) of all Np_n_N levels were detected for Ap_3_G in a medium missing methionine and cysteine during 16 and 40 hours of growth.

In summary, we have developed and validated a method for the simultaneous detection and quantification of highly negatively charged signaling molecules using WAX extraction in conjunction with CE-MS. Applying this to *E. coli* under varying stress conditions has unveiled a first glimpse at an apparently highly complex signaling network that is far from being understood.

## Methods

### Bacterial growth

The *E. coli* K12 strain BW25113 was chosen as wild type strain and the corresponding *ΔrelA* (Kan^R^) strain from the Keio collection was purchased from Horizon Discovery. For reducing the risk of second site suppressors or revertants, the strains were streaked freshly from glycerol stock before each experiment and propagated as a liquid culture at 37 °C, 180 rpm agitation in a 1:5 liquid to vessel volume, unless otherwise stated. The main cultures were inoculated from overnight precultures using a ratio of 1 × 10^8^ cells/10 mL fresh prewarmed medium. The cell proliferation was monitored by regular optical density measurements.

For analysis of (p)ppGpp under non-stressed conditions, cells were grown in 0.4 % glucose-enriched MOPS medium^[15,58]^ the Neidhardt Supplementary Mixture, Complete was purchased from Formedium (Norfolk, England).

M63 medium supplemented with 2 % Glycerol as a carbon source and enriched with all 20 proteinogenic amino acids (0.1 mM each) was prepared according to Ausubel et. al.^[69]^. For inducing partial amino acid starvation, Methionine, Cysteine, or both were omitted from the amino acid mix, but pre-cultures were grown in the presence of 20 amino acids.

Sample acquisition for all experiments was performed by quick decanting a discrete volume of cell culture in prelabeled, prechilled tubes containing 25 M formic acid, diluting it to a final concentration of 1 M, followed by quick and thorough mixing to ensure pH-induced cell lysis and nucleotide fixation. Samples were then flash frozen in liquid nitrogen and stored at -80 °C until extraction. Once the sample was acquired, a subsequent immediate OD_600_ measurement was performed to assess the cell density of the cell suspension and to extrapolate the cell number in each sample.

### Extraction

Extractions were performed using the protocol of Varik et al.^[15]^ Lysed bacteria were thawed at 38 °C and spiked with internal heavy standards of [^15^N]_5_ (p)ppGpp (synthesized in our lab^[50]^) and commercially available [^13^C]_10_ ATP and GTP (Sigma Aldrich, used without further purification). Cell lysates were vortexed every 2 minutes during incubation for 30 minutes. Samples were diluted with NH_4_OAc (1 ml per 3.125 ml lysate, 50 mM, pH = 5.5) and centrifuged (10 min, 3220 g, 4 °C). In the automated weak anion extraction, using a GX-241 ASPEC from Gilson, cartridges from biotage (EVOLUTE® EXPRESS WAX, 100 mg/3 ml (Tabless), # 614-0010-BXG) were equilibrated with MeOH (1 ml) and NH_4_OAc (1 ml, 50 mM, pH = 4.5). Samples were loaded onto the cartridge were washed with NH_4_OAc (1 ml, 50 mM, pH = 4.5) and MeOH (1 ml) and eluted with MeOH:ddH_2_O:NH_4_OH buffer (20:70:10, 2 × 750 µl). After dilution with ddH_2_O (2 ml) samples were lyophilized overnight. Residuals were dissolved in ddH_2_O (100 µl) and the solvent was removed via vacuum concentration (5 h, RT, V-AQ). Samples were dissolved in ddH_2_O (30 µl) and stored at -20 °C until preparation for measurement.

### CE measurements

Samples were measured on a commercially available capillary electrophoresis system coupled to mass spectrometry from Agilent. An Agilent 7100 CE system was coupled via an Agilent jet stream (AJS) electrospray ionization (ESI) source and an Agilent liquid coaxial interface to mass spectrometry. An Agilent G6495C Agilent QQQ or an Agilent 6545 qTOF mass spectrometer was used for detection. Analyte solution was diluted via an Agilent 1200 isocratic LC pump and a 1:100 splitter resulted in a flow of 10 µl/min with ddH_2_O-isopropanol mixture (1:1 v-%), spiked with internal mass standards in Q-TOF measurements. In measurements a bare fused silica capillary (100 cm length, 50 μm internal diameter) was used. Activation of the capillary was performed with 1 M NaOH and water for 10 min before first measurement. Washing of the capillary with ddH_2_O (300 s) and separation buffer ammonium acetate (35 mM, pH = 9.75, 300 s) were done before each measurement. Samples were diluted with ddH_2_O in a 1:1 ratio and injected by applying pressure (100 mbar, 20 s). Separation was achieved using a stable current of 22 µA via applying a voltage of +30 kV.

### QqQ measurements

QqQ measurements were performed on an Agilent G6495C Agilent QQQ in negative ionization mode. Capillary voltage was set to -2000 V and nozzle voltage to 2000 V. Pressure RF was between 60 to 90 V and the nebulizer gas was set to 8 psi, a temperature of 150 °C and a flow of 11 l/min. Sheath gas settings were a temperature of 175 °C and a flow of 8 l/min. Peaks were assigned via internal heavy standards and MS/MS transitions (see table 6). Agilent MassHunter Optimizer Automated MS Method Development Software (version 10.1) was used for determination of MS/MS transitions. Data acquisition was done with Agilent Mass Hunter Workstation LC/MS Data Acquisition for 6400 Series Triple Quadrupole (version 10.1) and analysis of the data was done with Agilent MassHunter Workstation Quantitative Analysis for QQQ (Quant-My-Way, version 10.1) and Agilent MassHunter Workstation Qualitative Analysis Navigator (version B.08.00).

### Q-TOF measurements

Q-TOF measurements were performed on an Agilent 6545 qTOF mass spectrometer in negative ionization mode. ESI spray current was set to 2.0 µA and the capillary voltage to -3500 V. Further MS settings were 100 V as fragmentor voltage, 65 V as skimmer voltage and 750 V as Oct RFV voltage. Drying gas was set to 250 °C with a flow of 3 l/min. The nebulizer gas had a pressure of 5 psi. Reference masses (TFA anion, [M-H]^-^, 112.9855 and HP-0921, [M-H+CH_3_COOH]^-^, 980.0163) were used for automatic recalibration of spectra with a rate of 1 spectra/min over a m/z range of 80-1700. A 10 ppm mass tolerance window of the corresponding theoretical masses of compounds was used to obtain extracted ion chromatograms (EICs). Data acquisition was done with Agilent Mass Hunter Workstation LC/MS Data Acquisition for 6200 Series TOF/6500 series Q-TOF (version 10.1) and analysis of the data was done with Agilent MassHunter Workstation Quantitative Analysis for Q-TOF (Quant-My-Way, version 10.1) and Agilent MassHunter Workstation Qualitative Analysis Navigator (version B.08.00). Assignment of peaks were achieved by accurate mass and identical migration time compared to heavy standards, if available.

## Acknowledgements

This project has received funding from the European Research Council (ERC) under the European Union’s Horizon 2020 research and innovation program (grant agreement no. 864246 to H.J.J.). H.J.J. acknowledges funding from Volkswagen Foundation (VW Momentum Grant 98604). This study was supported by the Deutsche Forschungsgemeinschaft (DFG) under Germany’s Excellence Strategy (CIBSS, EXC-2189, Project ID 390939984, to H.J.J.)

