## Supplemental Information for "Simultaneous Capillary Electrophoresis - Mass Spectrometry profiling of (p)ppGpp and dinucleoside polyphosphates reveals oscillatory (p)ppGpp dynamics and a potential synchronisation between alarmones"

### Results

#### Levels of ppGpp during growth

Next to the two growth curves displayed in the main publication, two other repeats were extracted and measured during exponential phase in MOPS medium. All ppGpp levels during growth showing a first peak in the middle of exponential phase and later a second increase (both increases marked with green arrows). Moreover, fluctuated levels were detected and are also shown in figure S1. As *E. coli* grew with different growth rates, turning points in $OD_{600}$ measurements were determined and time scale was accordingly shifted. With this calculation, first spikes of ppGpp levels were detected at the same time. Different absolute levels of ppGpp could be explained with mutations on the relA or SpoT allele, which appear frequently in K-12 stains[1].

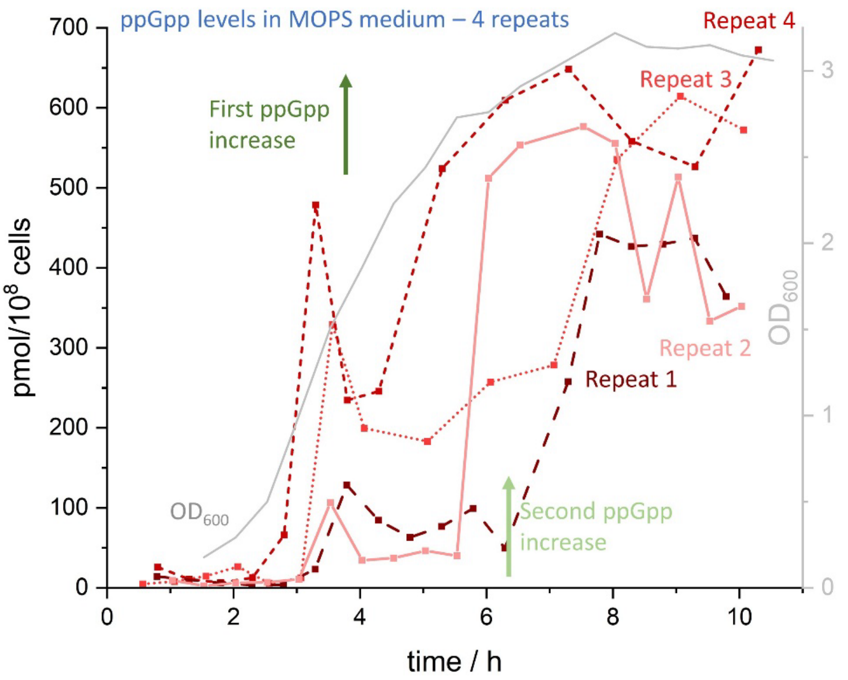


Figure S 1: Four repeats of ppGpp levels during growth. Increases of ppGpp are marked with green arrows (first increase: dark green, second increase: light green). Repeat 1 is represented by a dotted line with bigger lines and in dark red, repeat 2 is represented by a straight line and light red. Repeats 3 and 4 are shown in a small dotted line in lighter red and as small lines in darker red, respectively. In grey, $OD_{600}$ of one experiment is displayed. For better comparison, the time scale was adjusted so that the turning point in $OD_{600}$ measurements were at the same time. Exact values are provided in an xls source file.

After 18 hours of growth ppGpp levels in M63 medium supplemented with glucose and 18 amino acids (methionine and cysteine missing), ppGpp levels decreasing (12.2-fold, figure S2). Oscillations like described in exponential growth were not detected. This could be because of too large measurement intervals as also seen in other literature[2].

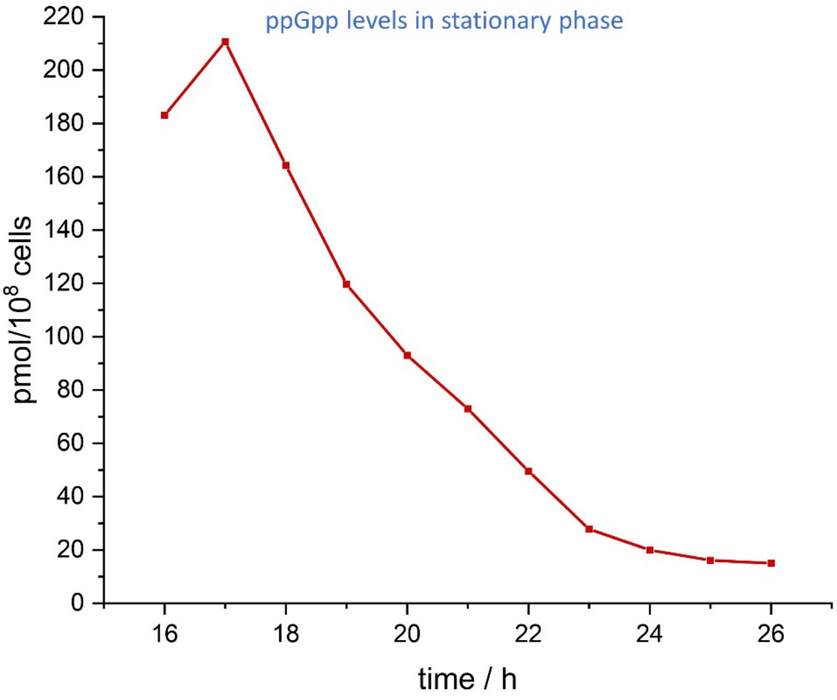


Figure S 2: Decreasing ppGpp levels after 16 hours of growth in M63 medium. ppGpp is displayed in red.

In further growth, ppGpp level still decreased (20 AA: 2.4-fold, 19 AA-Met: 2.3-fold, 19 AA-Cys: 2.7-fold, 18 AA-Met-Cys: 1.3-fold), as shown in figure S3. These decreases were independent of the amino acid supplementation in the growth medium of bacteria.

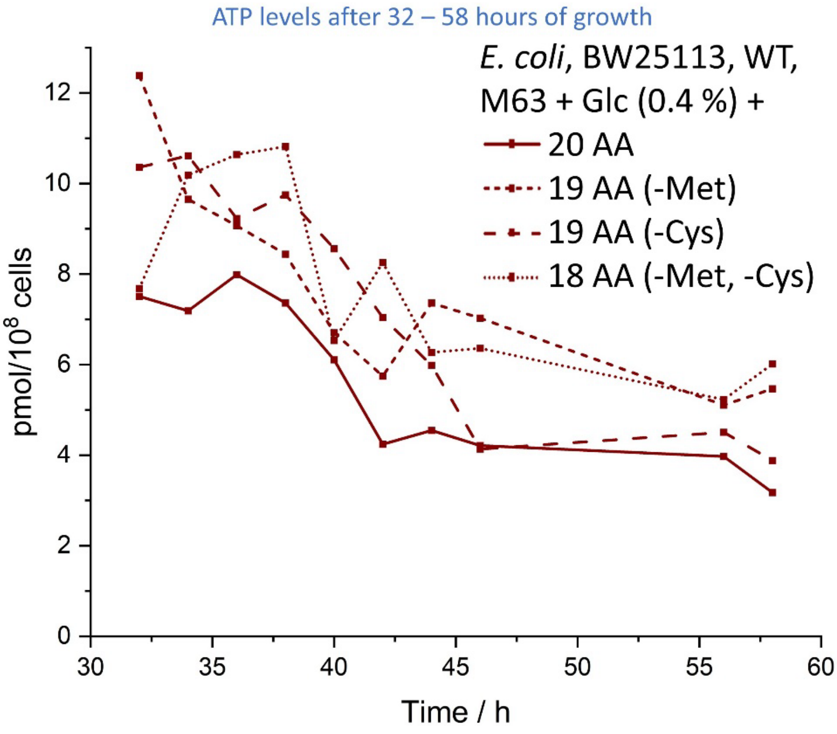


Figure S 3: Decreasing levels of ppGpp after 32 to 58 hours of growth.

### Levels of pppGpp during growth

In figure S4 all determined levels of pppGpp in MOPS medium are shown. The characteristic ppGpp pattern of a first peak in ppGpp levels followed by a decrease and a second increase, is for pppGpp not as profound as for ppGpp. Nevertheless, the mathematically predicted fluctuations[3] of this second messenger can be detected. Like for ppGpp levels, growth shifts of E. coli were corrected by the turning point of $OD_{600}$ measurements.

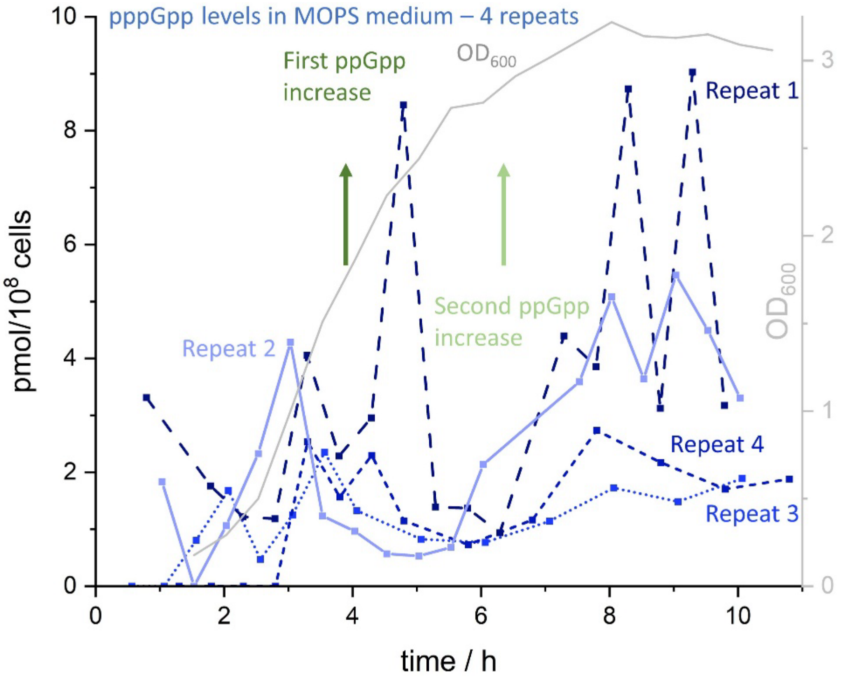


Figure S 4: Four repeats of pppGpp levels during growth. Increases of ppGpp are marked with green arrows (first increase: dark green, second increase: light green). Repeat 1 is represented by a dotted line with bigger lines and in dark blue, repeat 2 is represented by a straight line and light blue. Repeats 3 and 4 are shown in a small dotted line in lighter blue and as small lines in darker blue, respectively. In grey, $OD_{600}$ of one experiment is displayed. For better comparison, time scale was shifted so that the turning point in $OD_{600}$ measurements were at the same time. Exact values are provided in an xls source file.

### $Np_nN$ levels after growth of 32 to 58 hours in medium supplemented with 20 amino acids

For comparison of amino acid starvation conditions and adequate growth conditions, *E. coli* grew in M63 medium supplemented with all 20 amino acids and glucose. In this condition 8 different $Np_nNs$ were found. The missing dinucleoside polyphosphate in comparison with amino acid starvation was $Cp_4U$. As $Cp_4U$ was also the less abundant $Np_nN$ in *E. coli* grew in amino acid starvation condition, this $Np_nN$ was probably below the detection limit. Time changes of all detected dinucleoside polyphosphates are shown in figure S5.

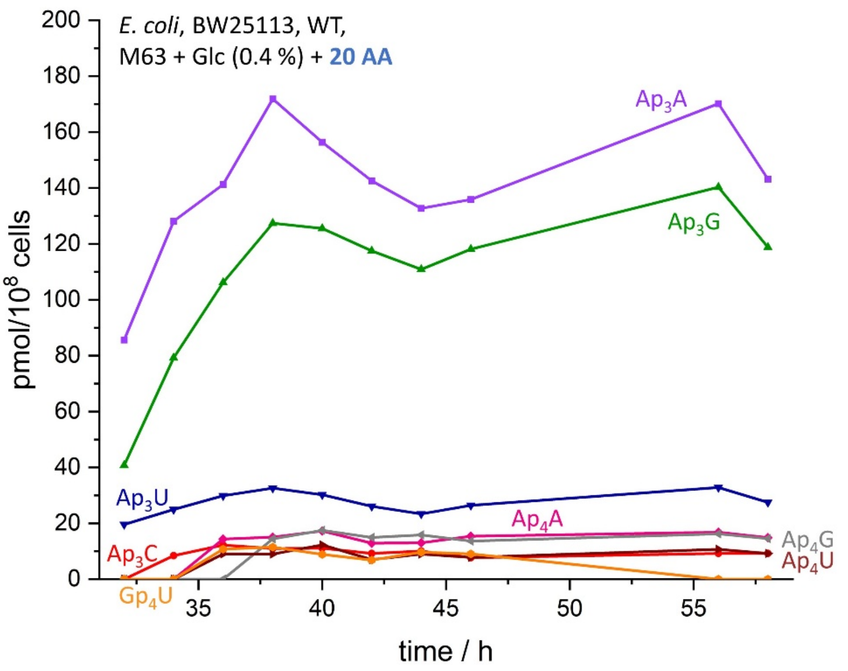


Figure S 5: $Np_nN$ levels after 32 to 58 hours of growth with addition of all 20 amino acids in M63 medium. All media contained M63 and glucose (0.4 %). Represented are $Ap_3A$ in purple, $Ap_3C$ in red, $Ap_3G$ in green, $Ap_3U$ in blue, $Ap_4A$ in pink, $Ap_4G$ in grey, $Ap_4U$ in brown, $Gp_4U$ in orange. Exact values are provided in an xls source file.

After extraction of 40 × $10^8$ *E. coli* (1.4-fold more material compared to $Np_nN$ level determination after 32 – 58 h of growth) eleven different dinucleoside polyphosphates were found. In figure S6 relative concentrations of all detected $Np_nNs$ are shown. Bacteria (BW25113) grew in M63 medium supplemented with glucose (0.4%) and 18 amino acids (- methionine and cysteine).

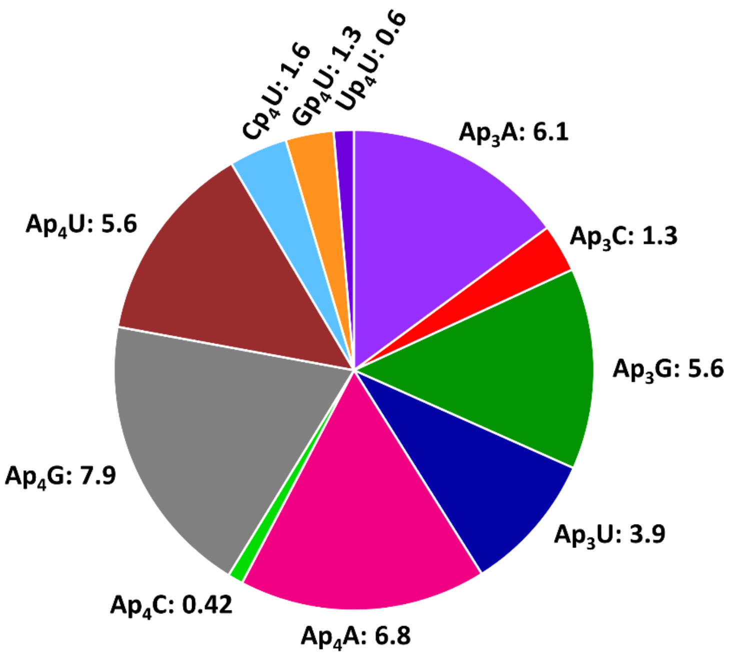


Figure S 6: Levels of $Np_nN$ after 24 hours of growth in M63 medium supplemented with 18 amino acids (- methionine and cysteine) and glucose. Light green field: $Ap_4C$ and a level of 0.4 pmol/$10^8$ cells. All levels are indicated in pmol/$10^8$ cells and fields sizes are relative to $Np_nN$ content in sample. $Np_nNs$ are depicted in the same colors like in Figure S5).

### Levels of ATP during growth

ATP levels in exponentially growing *E. coli* were determined in four different biological repeats in MOPS medium and are shown in figure S6. Small overall decreases (1 – 2.5-fold) can be detected. The same oscillations described for the levels of magic spots can be also detected in ATP levels.

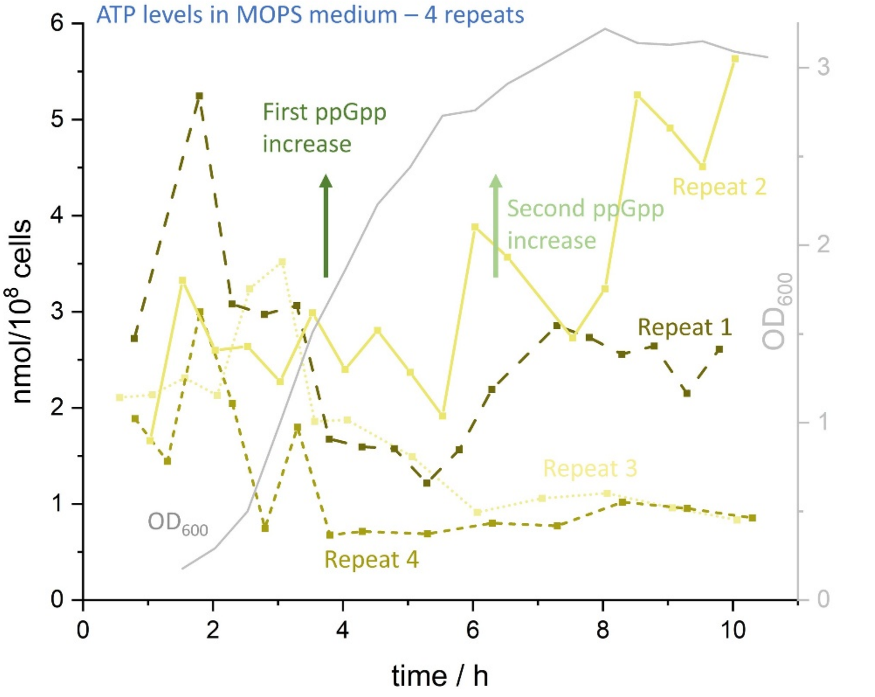


Figure S 7: Four repeats of ATP levels during growth. Increases of ppGpp are marked with green arrows (first increase: dark green, second increase in light green). Repeat 1 is represented by a dotted line with bigger lines and in dark yellow, repeat 2 is represented by a straight line and light yellow. Repeats 3 and 4 are shown in a small dotted line in lighter yellow and as small lines in darker yellow, respectively. In grey, $OD_{600}$ of one experiment is displayed. For better comparison, time scale was adjusted so that the turning point in $OD_{600}$ measurements were at the same time.

Next to $Np_nN$ and ppGpp levels, also ATP levels after 16 hours of growth in M63 medium supplemented with glucose and 18 amino acids (missing methionine and cysteine) were measured and are displayed in figure S7. Interestingly, ATP levels increase 3.1-fold, although $Np_nN$ levels are increasing in this growth period and ATP is one precursor of $Ap_4Ns$.

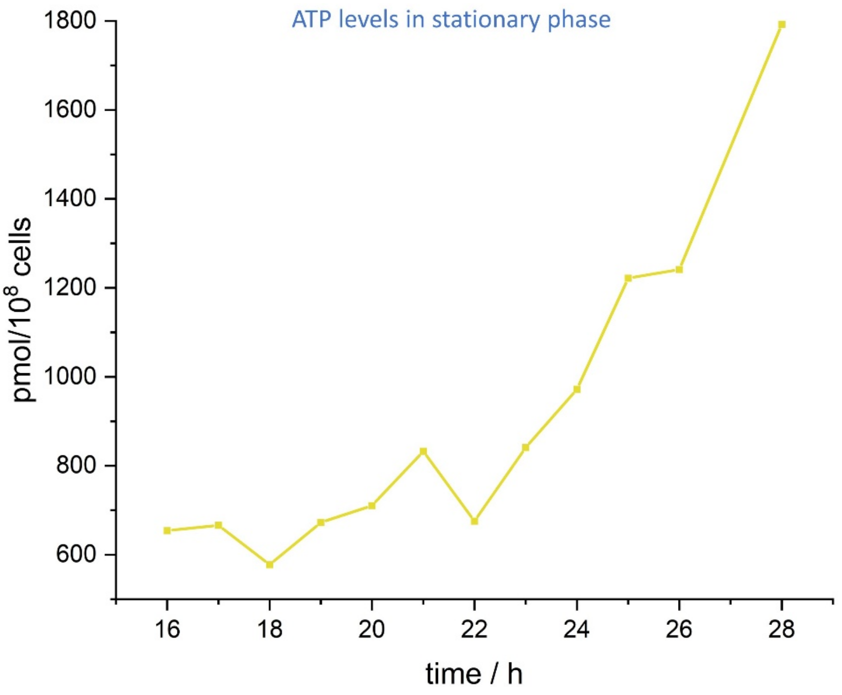


Figure S 8: Increasing ATP levels in stationary phase. ATP levels are shown in yellow.

In the last measurement interval presented here, ATP levels were also determined. ATP levels decreased during this time interval (20 AA: 51.8-fold, 19 AA-Met: 33.1-fold, 19 AA-Cys: 32.1-fold, 18 AA-Met-Cys: 57.6-fold) independent of the amino acid supplement as shown in figure S8.

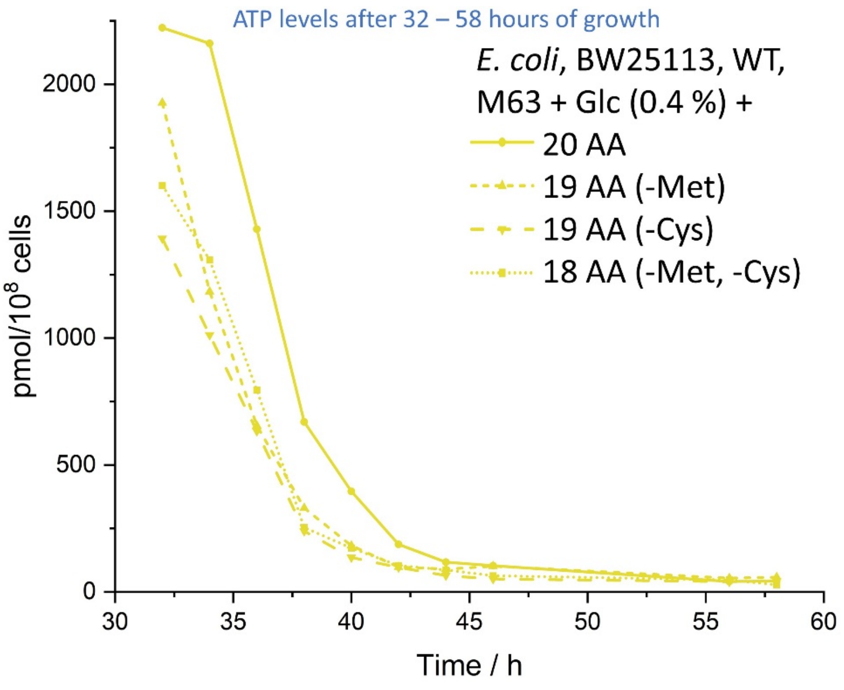


Figure S 9: Decreasing ATP levels after growth until 32 - 58 hours. ATP levels are displayed in yellow.

### ADP levels during growth

One precursor of $Ap_3Ns$ is ADP[4]. As $Ap_3G$ is one member of this group of molecules and increases more than 300-fold during this time, ADP levels were determined. As shown in figure S11, ADP levels are stable in this time period, but show fluctuations like observed for (magic spot) nucleotides.

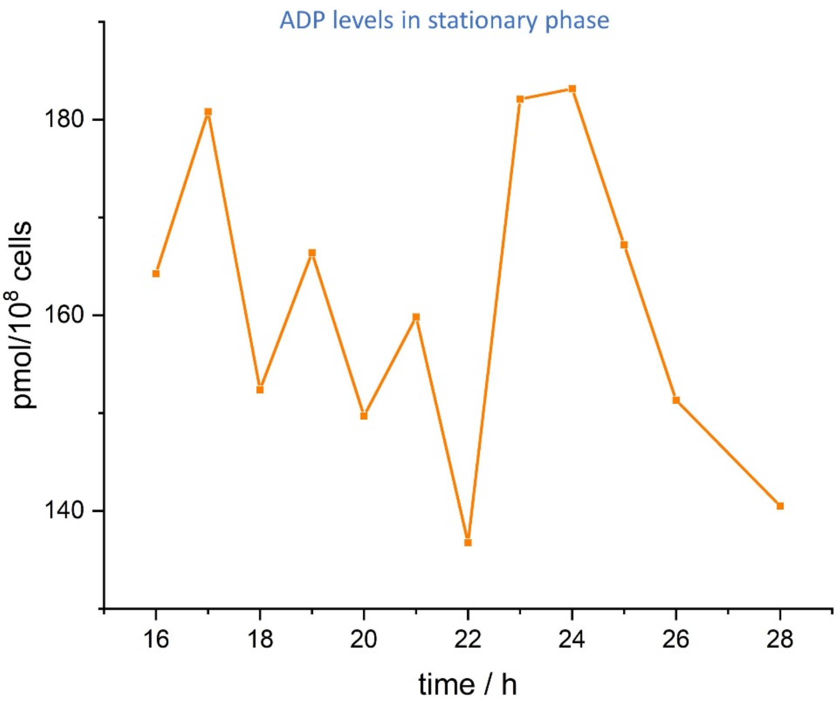


Figure S 10: ADP levels with stable but oscillating levels after 16 hours of growth. ADP levels are shown in orange.

ADP levels were also determined during growth between 32 – 58 hours of growth and are displayed in figure S12. The same decreases like detected in ATP are displayed in ADP levels (20 AA: 7.2-fold, 19 AA-Met: 2.7-fold, 19 AA-Cys: 6.3-fold, 18 AA-Met-Cys: 4.5-fold). These decreases were also independent of the amino acid supplementation of the medium.

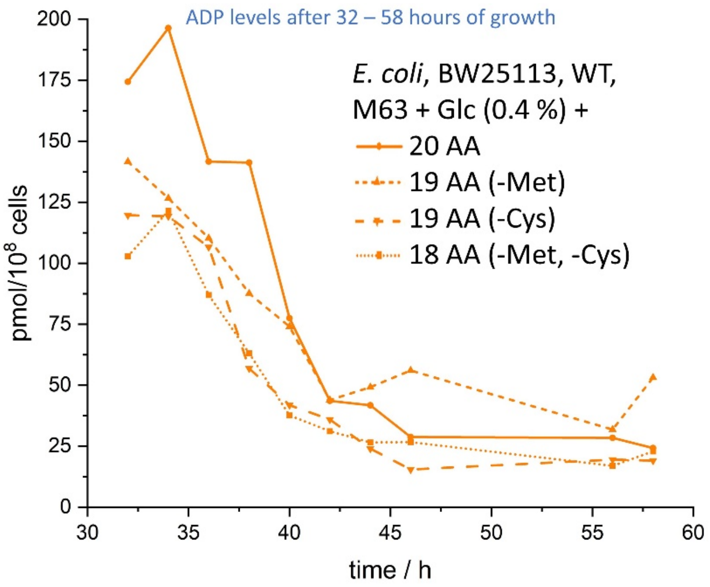


Figure S 11: Starvation independent decreases in ADP levels of different amino acid supplementation after growth of 32 - 58 hours.

### Methods

#### Limit of detection (LOD) and limit of quantification (LOQ) of $Ap_3A$

$^{18}O_1$-$Ap_3A$ were spiked in extracted bacterial sample (30 × $10^8$ cells). Measurements of triplicates were performed on a CE-QqQ system. A definition of peaks higher than a signal to noise ratio (S/N) of 3:1 for LOD and 10:1 for LOQ was used. The LOD was determined as 34 nM and the LOQ as 1 µM. Chromatograms of the LOD (figure S13) and LOQ (figure S14) are displayed below. Plotted are also the automated and manual noise selection, as the automated noise selection was pretentious. Calculations were performed by the data analysis software.

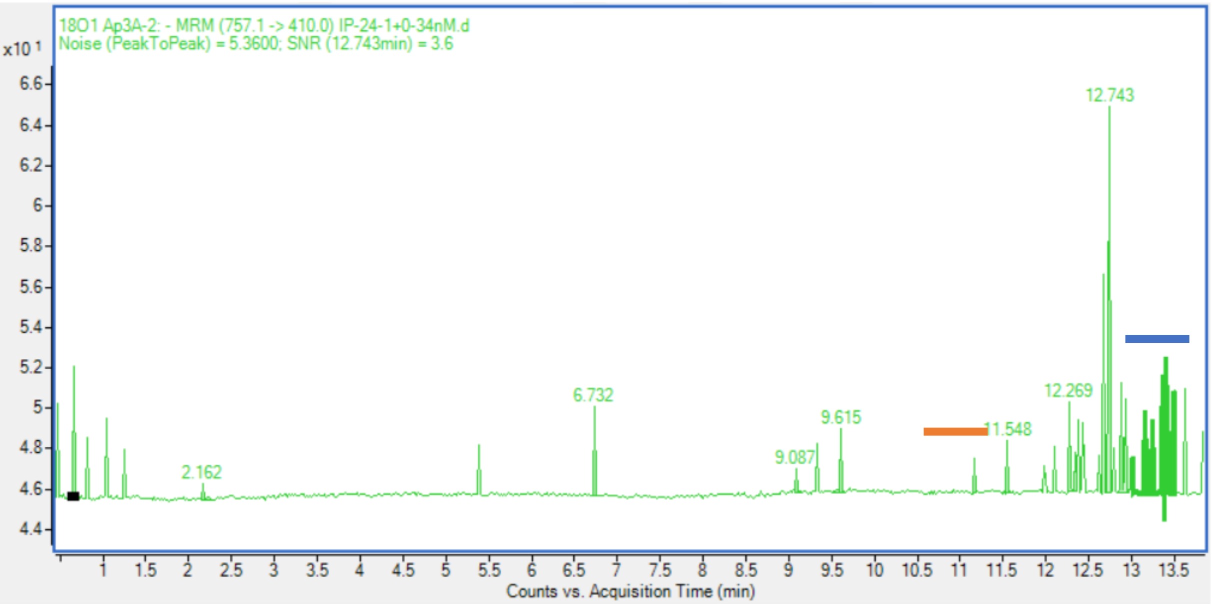


Figure S 12: Determination of LOD. Plotted are the automated (orange) and manual (blue) noise description.

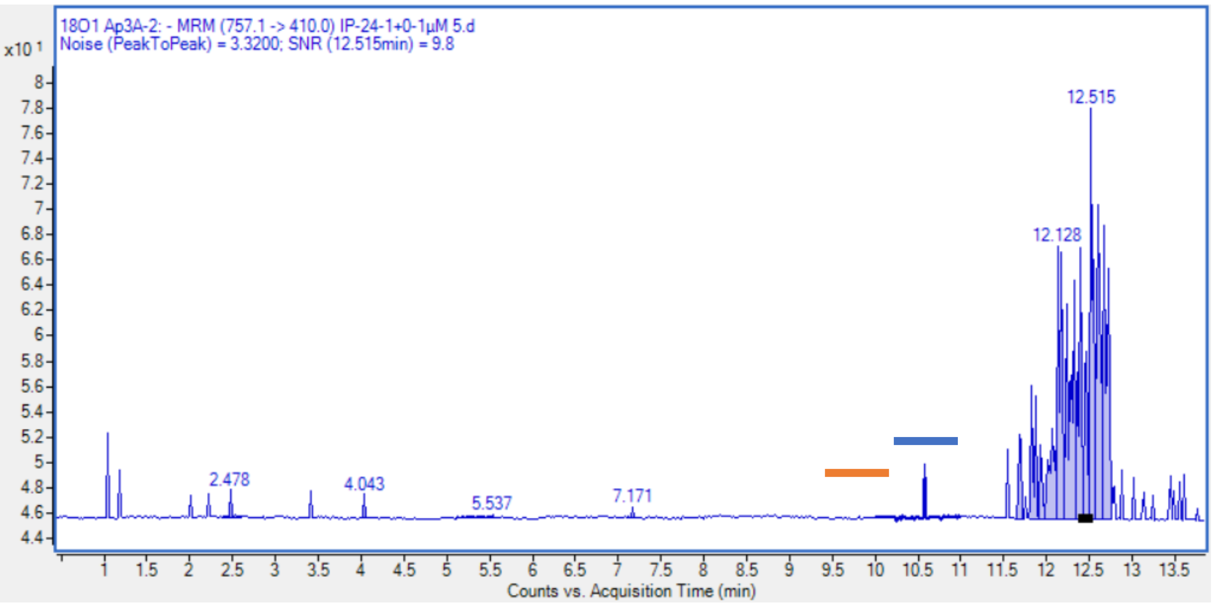


Figure S 13: Determination of LOQ. Plotted are the automated (orange) and manual (blue) noise description.

### Limit of detection (LOD) and limit of quantification (LOQ) of $Ap_4A$

Biological samples treated like samples in measurements but without detectable $Ap_4A$ were spiked with a $^{12}C$-$Ap_4A$ standard in different concentrations (LOD: 2 µM, 3 µM, 4 µM, 5 µM, 7 µM, 8 µM; LOQ: 7 µM, 8 µM, 10 µM, 11 µM, 12 µM). Abundances of $Ap_4A$ peaks were analyzed. Counts vs concentrations were displayed and a linear regression of triplicates were calculated (one exemplified shown in figure S14 (LOD) and S15 (LOQ), respectively). LOD and LOQ were calculated using the standard deviation of the Y-intercept (SD), the slope (m) of the linear regression (OriginPro 2025, Data analysis and Graphing Software, 2025b) and the following equation (SE1 (LOD)[5] and SE2 (LOQ)[6]):

$$LOD = \frac{3.3 \cdot SD}{m} \qquad \text{(SE1)}$$

$$LOQ = \frac{10 \cdot SD}{m} \qquad \text{(SE2)}$$

Errors were calculated using the standard deviation of the determined LOD and LOQ for each experiment. The LOD were calculated as 2.0 ± 0.3 µM and LOQ as 6.8 ± 1.2 µM.

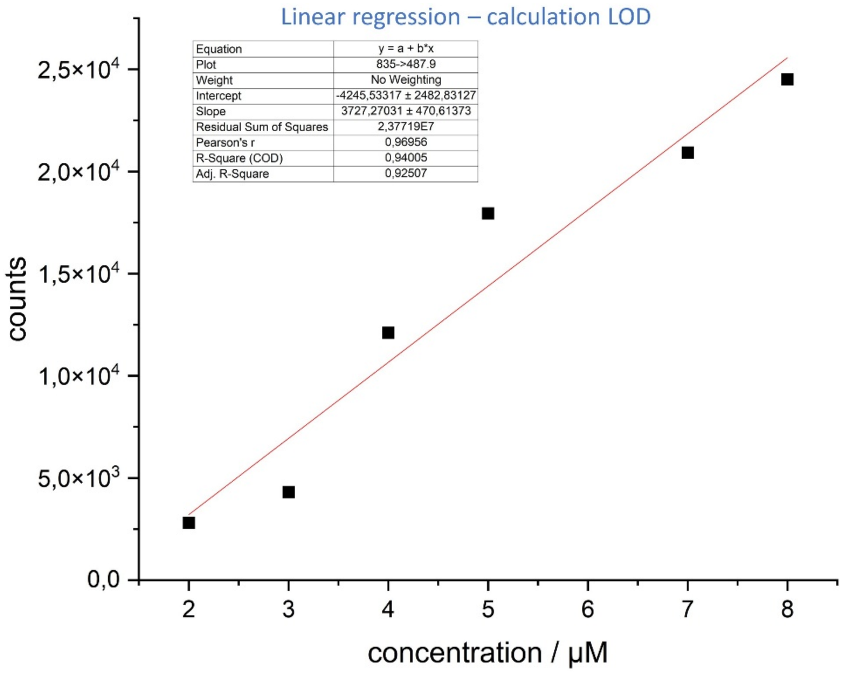


Figure S 14: One exemplified linear regression used for the calculation of the limit of detection.

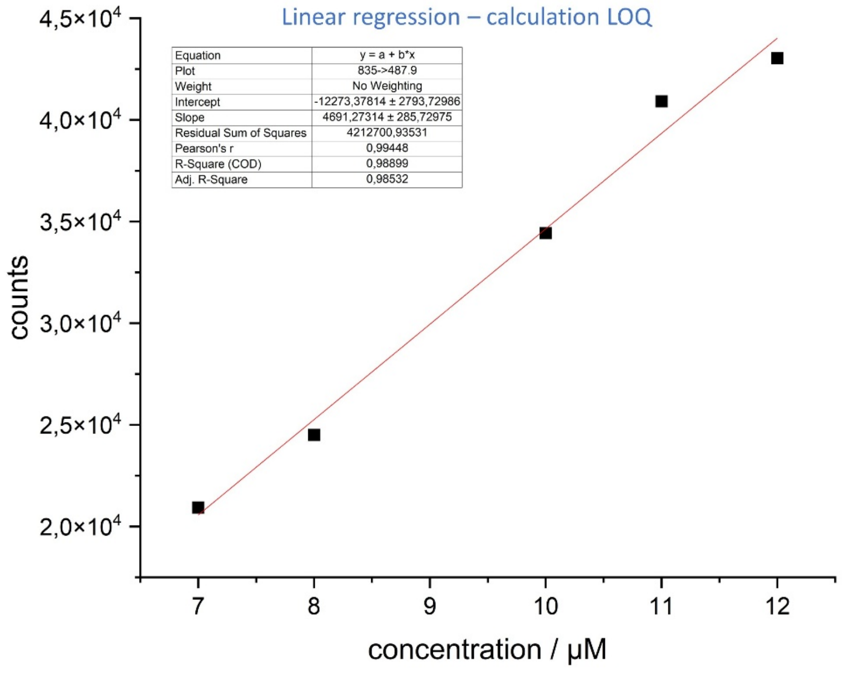


Figure S 15: One exemplified linear regression used for the determination of the limit of quantification.

### Recovery of $Ap_3A$ and $Ap_4A$

Recoveries of $Ap_3A$ and $Ap_4A$ were determined using the described, slightly modified method of Ihara et al.[7] and Bartoli et al.[8]. Commercially available standards of $Ap_3A$ and $Ap_4A$ were dissolved in water. Two different concentrations of the aqueous solutions were incubated for 30 minutes on ice after addition of formic acid to a final concentration of 1 M. No further changes of the protocol described before were made.

After extraction of the standard solution, abundances of CE-QqQ measurements were compared with unextracted solutions. The recoveries of two different concentrations were determined, because Cohen et al.[9] reported different recoveries depending on the concentration of the tested nucleotides. Results were calculated using triplicates of independent extracted samples.

### Calculated and found HRMS of $Np_nNs$

Many of the detected dinucleoside polyphosphates are not commercially or synthetically available and were therefore only identified by high resolution mass (HRMS). Important for the identification of a molecule is the mass variance, which has to be smaller than 5 ppm. As shown in table S1, all of the calculated mass accuracies are smaller than the required value of 5 ppm. Nevertheless, the structural identification is not clear with the HRMS. Measurements using the more sensitive CE-QqQ were not possible, because without a pure standard of each $Np_nN$ the knowledge about the fragmentation pattern is missing.

Table S1: Calculated and found high resolution masses of all detected dinucleoside polyphosphates. Moreover, variances of detection and calculations as well as number of MS scans are displayed.

| $Np_nN$ | HRMS calculated | HRMS found | Variance | Mass accuracy / ppm | # scans |
|---|---|---|---|---|---|
| **$Ap_3A$** | 755.0747 | 755.0734 | 0.0013 | 1.7 | 39 |
| **$Ap_3C$** | 731.0634 | 731.0643 | 0.0011 | 1.2 | 38 |
| **$Ap_3G$** | 771.0696 | 771.0688 | 0.0008 | 1.0 | 38 |
| **$Ap_3U$** | 732.0474 | 732.0464 | 0.0010 | 1.4 | 38 |
| **$Ap_4A$** | 417.0169 | 417.0168 | 0.0001 | 0.2 | 44 |
| **$Ap_4C$** | 413.0087 | 413.0101 | 0.0014 | 3.4 | 22 |
| **$Ap_4G$** | 425.0143 | 425.0141 | 0.0002 | 0.5 | 45 |
| **$Ap_4U$** | 405.5032 | 405.5030 | 0.0002 | 0.5 | 32 |
| **$Cp_4U$** | 393.4976 | 393.4972 | 0.0004 | 1.0 | 32 |
| **$Gp_4U$** | 413.5007 | 413.5002 | 0.0005 | 1.2 | 38 |
| **$Up_4U$** | 393.9896 | 393.9903 | 0.0007 | 1.8 | 23 |

### $Ap_3A$ peak assignment

Molecules with the same chemical formula have the same mass. Nevertheless, the structure of these molecules can be different (structural isomers). The characterization of the structure can be done using NMR, fragmentation pattern or identical retention time on separation columns. Whereas, the first method needs pure compounds, the latter two are needing standards for comparison. As we wanted to know the retention time of $Np_nNs$ in CE runs and detected the same fragmentation pattern in QqQ measurements, we spiked one of our biological sample with heavy $^{18}O_1$-$Ap_3A$. The heavy standard was synthesized in our lab according to the published synthesis route[10]. The extracted CE run is shown in figure S17.

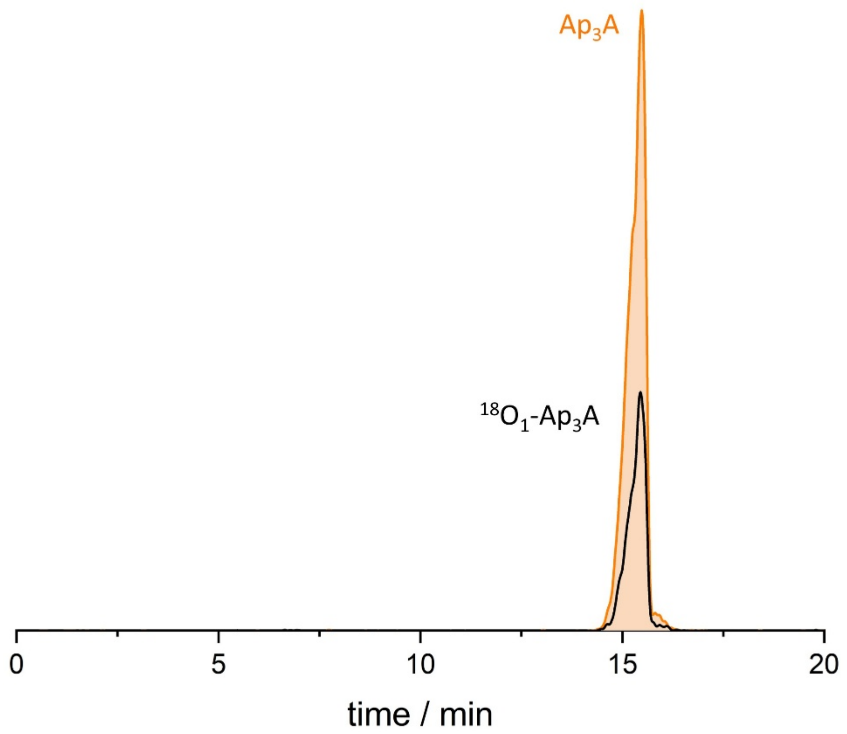


Figure S 16: $Ap_3A$ peak of sample and heavy $^{18}O_1$-$Ap_3A$ as internal standard. The peaks with same retention time confirm structural identification of $Ap_3A$.
